# Recruitment of mRNAs to P granules by gelation with intrinsically-disordered proteins

**DOI:** 10.1101/732057

**Authors:** Chih-Yung S. Lee, Andrea Putnam, Tu Lu, Shuaixin He, John Paul T. Ouyang, Geraldine Seydoux

**Affiliations:** HHMI and Dept. of Molecular Biology and Genetics, Johns Hopkins University School of Medicine, Baltimore MD USA; Dept. of Biophysics and Biophysical Chemistry, Johns Hopkins University School of Medicine, Baltimore MD USA; Google, 1600 Amphitheatre Parkway, Mountain View, CA, USA

**Author notes:** Embryonic P granules are super-assemblies of nanoscale RNA-protein gels.

## Abstract

Animals with germ plasm assemble cytoplasmic RNA granules (germ granules) that segregate with the embryonic germ lineage. How germ granules assemble and recruit RNA is not well understood. Here we characterize the assembly and RNA composition of the germ (P) granules of *C. elegans*. ∼500 maternal mRNAs are recruited into P granules by a sequence independent mechanism that favors mRNAs with low ribosome coverage. Translational activation correlates temporally with P granule exit for two mRNAs that code for germ cell fate regulators. mRNAs are recruited into the granules by MEG-3, an intrinsically disordered protein that condenses with RNA to form nanoscale gels. Our observations reveal parallels between germ granules and stress granules and suggest that cytoplasmic RNA granules are reversible super-assemblies of nanoscale RNA-protein gel condensates.

## Introduction

RNA granules are RNA/protein condensates that assemble in the cytoplasm or nucleoplasm of cells. Unlike other organelles, RNA granules are not bound by membranes and can assemble and disassemble within minutes in cells. The dynamic properties of RNA granules have suggested that these structures are liquid droplets that assemble by phase separation (Alberti et al., 2019; Boeynaems et al., 2018; Hyman et al., 2014; Lin et al., 2015; Mittag and Parker, 2018; Shin and Brangwynne, 2017; Weber and Brangwynne, 2012). This hypothesis has been supported by *in vitro* experiments reporting liquid-liquid phase separation of RNA-binding proteins containing intrinsically-disordered domains (IDRs) (Brady et al., 2017; Lin et al., 2015; Nott et al., 2015; Shin and Brangwynne, 2017). IDRs have been proposed to mediate labile protein-protein interactions that drive phase separation to form round droplets held together by surface tension (Banani et al., 2017; Boeynaems et al., 2018; Mittag and Parker, 2018; Shin and Brangwynne, 2017). Observations in cells, however, have revealed that cytoplasmic RNA granules often adopt complex, irregular morphologies, with non-uniformly distributed proteins and RNAs. For example, stress granules contain stable, protein dense, irregular “cores” surrounded by more labile and dilute “shells” (Cirillo et al., 2019; Jain et al., 2016; Wheeler et al., 2016; Woodruff et al., 2018). Similarly, the Balbiani body of zebrafish and *Xenopus* embryos is built around a non-dynamic matrix with heterogeneous zones of protein concentration (Boke et al., 2016; Roovers et al., 2018; Woodruff et al., 2018). mRNAs have also been reported to be non-homogeneously distributed in granules: for example, mRNAs in the germ granules of *Drosophila* sort into homotypic clusters that adopt characteristic positioning within each granule (Eagle et al., 2018; Little et al., 2015; Trcek et al., 2015; Van Treeck et al., 2018). The mechanisms that underlie the heterogeneous organization of RNA granules are not well understood, although both protein and RNA aggregation mechanisms have been suggested (Van Treeck and Parker, 2018).

P granules are the germ granules of *C. elegans* and a well-studied model for cytoplasmic RNA granules (Marnik and Updike, 2019; Seydoux, 2018; Strome, 2005). P granules are heterogenous assemblies that contain at least two phases with distinct dynamics: a liquid phase assembled by the RGG domain proteins PGL-1 and its paralog PGL-3 (Brangwynne et al., 2009; Hanazawa et al., 2011; Updike et al., 2011), and a gel phase, assembled by the intrinsically-disordered proteins MEG-3 and its paralog MEG-4 (Putnam et al., 2019). In *vitro* and *in vivo*, MEG-3 forms small, non-dynamic condensates (<500 nanometers) that associate with the surface of the larger PGL condensates (>500 nanometers). In the one-cell zygote, MEG condensates enrich in the posterior cytoplasm (germ plasm) where they recruit and stabilize PGL condensates inherited from the oocyte (Putnam et al., 2019; Smith et al., 2016). Preferential assembly and stabilization of P granules in germ plasm ensures their preferential inheritance by germline blastomeres (Fig. S1A). By analogy to other germ granules, P granules are thought to deliver mRNAs coding for germline determinants to the nascent germline (Strome, 2005). *In situ* hybridization experiments have shown that P granules contain polyadenylated mRNAs (Seydoux and Fire, 1994), but so far only one mRNA has been reported to localize to P granules in early embryos: *nos-2* (Subramaniam and Seydoux, 1999). *nos-2* codes for a homolog of the conserved germline determinant Nanos that specifies germ cell fate redundantly with *nos-1*, another Nanos homolog expressed later in development (Subramaniam and Seydoux, 1999). The mechanisms that recruit *nos-2* and possibly other maternal mRNAs to P granules are not known.

In this study, we use immunoprecipitation, genetic and *in situ* hybridization experiments to characterize the P granule transcriptome. We find that mRNAs are recruited into P granules by the MEG phase in a sequence-independent manner that favors long RNAs with low ribosome occupancy. Using *in vitro* reconstitution experiments, we show that MEG-3 interacts electrostatically with RNA to form non-dynamic gel-like condensates. The condensates cluster on the surface of PGL droplets to form reversible super-assemblies. Our findings reveal parallels between stress granules and P granules and suggest a revised model for RNA granule assembly.

## RESULTS

### Identification of P granule mRNAs by immunoprecipitation with MEG-3

To identify RNAs that associate with P granules *in vivo*, we performed Individual-nucleotide resolution UV crosslinking and immunoprecipitation (iCLIP) experiments (Huppertz et al., 2014) on MEG-3 and PGL-1 proteins tagged at each locus with GFP. We chose these two proteins because MEG-3 and PGL-1 are essential (with their respective paralogs MEG-4 and PGL-3) to assemble the gel (MEG) and liquid (PGL) phases of P granules (Hanazawa et al., 2011; Updike et al., 2011). Early embryos (1 to 100-cell stage) were exposed to ultraviolet light, lysed, and the cross-linked protein/RNA complexes were immunoprecipitated using an anti-GFP antibody (Fig. S1B). As a control, we also used embryos expressing GFP alone. The GFP immunoprecipitates were washed stringently, lightly treated with RNAse to trim the bound RNAs, extracted for RNA and deep-sequenced. Sequencing reads were mapped back to the *C. elegans* genome (ws235) and used to determine read counts per locus. Read counts obtained in the control GFP iCLIP were used to define background threshold. We identified 657 transcripts that were reproducibly recovered above the GFP background threshold in two independent MEG-3::GFP iCLIP experiments (“MEG-3-bound transcripts”; Fig. 1A, Fig. S1C and Table S1). In contrast, we identified only 18 transcripts above background in the two PGL-1::GFP iCLIPs, despite abundant PGL-1::GFP protein in the immunoprecipitates (Fig. 1A, Fig.S1B and Table S1). 15/18 of the PGL-1-bound transcripts were also in the MEG-3-bound list (Table S1). For each MEG-3-bound transcript, we compared the average normalized read count (RPKM) across the two iCLIPs to transcript abundance in embryos or in P blastomeres and found no strong correlation, suggesting that MEG-3 binds to a specific sub-set of mRNAs (Fig. 1B, Fig. S1D). Among the MEG-3-bound transcripts was *nos-2*, the only previously reported embryonic P granule transcript (Fig. 1B). To determine whether other MEG-3-bound transcripts localize to P granules, we used single molecule fluorescent *in situ* hybridization (smFISH) (Raj et al., 2008) to examine the distribution of various transcripts in embryos. We initially analyzed 9 transcripts: nos-2, 6 other transcripts in the MEG-3-bound list, and 2 transcripts recovered in the MEG-3 iCLIPs that did not meet the GFP background cut off. For each transcript, we determined the average granule size in the P2 blastomere and compared that to the average raw read count in the MEG-3::GFP iCLIPs and observed a strong correlation (R = 0.92) (Fig. 1C). In this analysis, *nos-2* clustered with two other genes also in the MEG-3-bound list (*F35G2.1 and F35C11.5*). Extrapolating from this correlation, we predicted that transcripts that ranked higher than the *nos-2* cluster in the MEG-3-bound list should all localize to P granules. In total, we examined 18 transcripts among this 492-gene set and found that, as predicted, all localized robustly to P granules (Fig. 1D, Fig. S2,S3). We also examined 7 transcripts that ranked below the *nos-2* cluster and found none that localized to P granules in all P blastomeres, although we observed occasional localization to smaller granules (Fig. S2). Finally, we examined 6 genes that were not recovered reproducibly in the iCLIPs and had above average expression in P blastomeres (RPKM=7). We found none that localized to P granules (Fig. 1D, Fig. S2). We conclude that ranking at or above the *nos-2* cluster in the MEG-3-bound list is a stringent metric for predicting transcripts with robust P granule localization. We designate this 492-gene set as “P granule transcripts”. This gene list corresponds to the top 75% of the 657 MEG-3-bound transcripts, 11% of the 1347 transcripts enriched in P blastomeres(148/1347) (Lee et al., 2017), and ∼3 % of the 15,345 transcripts expressed in early embryos.

**Figure 1.**
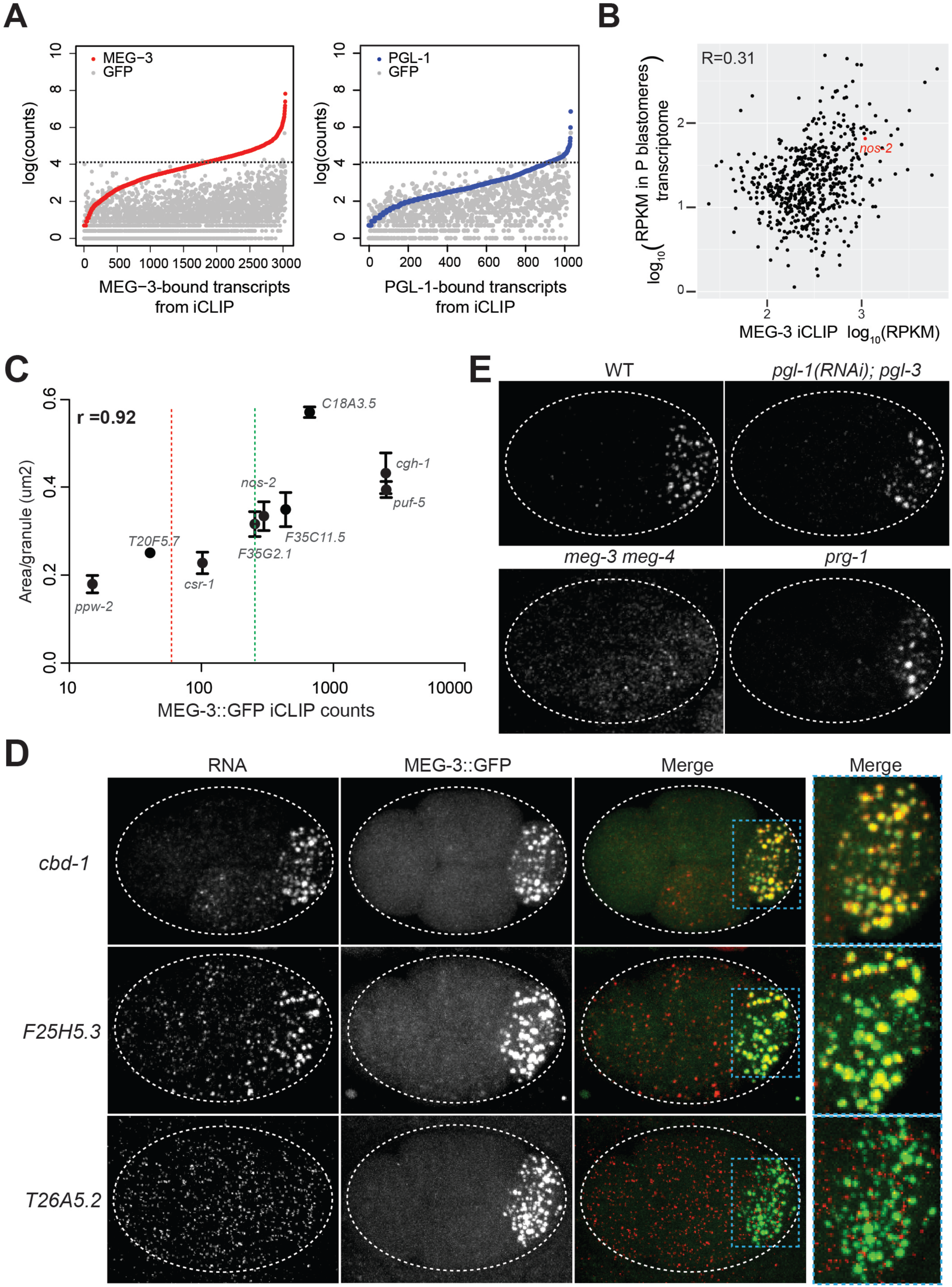
mRNAs are recruited into P granules by the MEG phase. (A) Graphs showing log transformed average read counts (Y axis) from two MEG-3::GFP (red), PGL-1::GFP (blue) and GFP (grey) iCLIP experiments. Genes are arranged along X axis based on the ascending log transformed read counts in the MEG-3::GFP or PGL-1::GFP iCLIP experiments (average of two experiments). Gray dots represent the read counts for the corresponding rank-ordered gene in the GFP iCLIP. The stippled line denotes the GFP background threshold (read counts =60) above which transcripts were considered true positives (657 transcripts in the MEG-3::GFP iCLIPs and 18 transcripts in the PGL-1::GFP iCLIPs). (B) Graph showing 657 MEG-3-bound transcripts (black dots) with respect to read counts in the MEG-3::GFP iCLIPs (average FPKM of two replicates, X axis) versus transcript abundance in P blastomeres (Y axis). R is the Pearson correlation coefficient. *nos-2* is highlighted in red. See Fig. S1D for graph comparing the same X axis versus transcript abundance in embryos. (C) Graph showing average read counts in MEG-3::GFP iCLIPs versus average RNA cluster size as measured from smFISH signal for 9 genes. R is the Spearman correlation coefficient. Red stippled line denotes threshold for MEG-3::GFP-bound mRNAs as defined in A. Green stippled line denotes threshold for P granule mRNAs (see text). (D) Photomicrographs of embryos expressing MEG-3::GFP hybridized with single molecule fluorescence (smFISH) probes as indicated. *cbd-1* and *F25H5.3* are examples of transcripts localizing to P granules (MEG-3::GFP), as shown at higher magnification in the right-most panels. *T26A5.2* is an example of a transcript that does not enrich in P granules. See Fig. S2 and Fig. S3 for additional examples. (E) Photomicrographs of 4-cell embryos of the indicated genotypes hybridized with smFISH probes against the *nos-2* transcript. All genotypes show localization of *nos-2* to P granules in the P_2_ blastomere, except for *meg-3 meg-4*. Note that *nos-2* is a maternal mRNA that is rapidly degraded in somatic blastomeres (two anterior cells) and therefore present at higher levels in P blastomeres and their newly born sister blastomeres (two posterior cells).

For all 18 P granule transcripts analyzed by smFISH, localization to P granules was inefficient with many molecules also found diffusely in the cytoplasm (Fig. 1D and Fig. S2, S3). We calculated the number of molecules in P granules and cytoplasm for two transcripts among the top 5 ranked in the MEG-3-bound list. We found that only 21 ±3 % *puf-5* and 34±3% *Y51F10.2* molecules localized to P granules in the P_2_ blastomere. Because P granules occupy only a small portion of the P_2_ cell volume (5.9±2 %), this enrichment corresponds to a ∼6 fold increase in concentration over the cytoplasm.

The iCLIP results suggest that mRNAs are recruited to P granules as part of the MEG phase rather than the PGL phase. Consistent with this hypothesis, *nos-2* mRNA still localized to granules in embryos lacking *pgl-1* and *pgl-3*, but not in embryos lacking *meg-3* and *meg-4* (Fig. 1E). We obtained similar results in *in situ* hybridization experiments against poly-A RNA and against *Y51F10.2*, one of the transcripts identified in both the MEG-3-bound and PGL-1-bound lists (Fig. S1E, Table S1). In *Drosophila*, mRNAs have been proposed to be recruited to germ granules via interaction with piRNAs complexed with the PIWI-class Argonaute Aubergine (Vourekas et al., 2016). We found that *nos-2* and *Y51F10.2* still localized to P granules in embryos lacking the PIWI-class Argonaute PRG-1 (Fig. 1E and Fig. S1E). We conclude that mRNA recruitment to embryonic P granules depends on MEG-3 and MEG-4 and does not require PGL-1 and PGL-3 or the Argonaute PRG-1.

### Characterization of P granule mRNAs: sequence non-specific preference for low ribosome-occupancy mRNAs

The majority of reads in the MEG-3::GFP iCLIPs mapped to protein-coding mRNA transcripts (Fig. S4A). The iCLIP protocol yields short (∼10-30bp) reads that correspond to sequences cross-linked to MEG-3::GFP (“footprints”) (Huppertz et al., 2014). Metagene analysis of the MEG-3::GFP footprints revealed that MEG-3 binds transcripts throughout the coding and UTR regions, with a 2X preference for 3’UTR sequences (Fig. 2A, Fig. S4A). We analyzed the MEG-3 footprints for possible motifs and found no evidence for any sequence preference (Methods). MEG-3-bound transcripts tended to be longer on average than other embryonic transcripts (Fig. 2B) and were enriched for transcripts known to be targeted by translational repressors expressed in oocytes, including OMA-1, GLD-1 and LIN-41 and CGH-1 (Boag et al., 2008; Scheckel et al., 2012; Tsukamoto et al., 2017) (Fig. S4B). Ribosome profiling experiments confirmed that MEG-3-bound transcripts are on average less protected by ribosomes than other embryonic transcripts (Fig. 2C, Fig. S4C for P granule transcripts). These observations suggested that MEG-3 recruits RNAs into P granules by a sequence non-specific mechanism that favors poorly translated mRNAs. This hypothesis predicts that, under conditions where translation is globally inhibited, previously cytoplasmic, well translated mRNAs should localize to P granules. We found that a brief incubation at 30°C (15 minutes heat shock) was sufficient to disassemble polysomes (McCormick and Penman, 1969; Shalgi et al., 2013) (Fig. S4D). We analyzed five non-P granule transcripts, chosen for their high ribosome occupancy under non-heat shock conditions, and remarkably found that all five accumulated in P granules after heat shock (Fig. 2D-E and Fig. S4E). Accumulation in P granules was observed in wild-type embryos, but not in embryos depleted of MEG proteins (Fig. S4E). We also noticed that MEG-3::GFP condensates become larger upon heat-shock. In zygotes, MEG-3 molecules exist in two states: a fast-diffusing, dilute pool in the cytoplasm and slow-diffusing condensed pool in P granules (Wu et al., 2019). Growth of MEG-3 granules under heat shock suggested that additional MEG-3 molecules condense in P granules under conditions when translation is repressed globally. To examine this further, we measured the size of MEG-3::GFP granules in embryos treated with drugs that block translation. For these experiments, we used *mex-5 mex-6* mutant embryos, which lack embryonic polarity and assemble MEG-3 granules throughout the cytoplasm (Smith et al., 2016). We found that treatment with puromycin, which causes ribosomes to dissociate from transcripts, increased the size of MEG-3::GFP granules by ∼4-fold, as also observed following heat-shock (Fig. 2F-H). In contrast, treatment with cycloheximide, which stalls ribosomes on transcripts, did not change the size of MEG-3::GFP granules (Fig. 2F-H). These findings parallel the divergent effect of puromycin and cycloheximide on the assembly of stress granules (Aulas et al., 2017; Kedersha et al., 2000), and suggest that MEG-3 recruits mRNAs to P granules by condensing in a sequence non-specific manner with ribosome-free mRNAs. We note, however, that not all poorly translated mRNAs associate with P granules with equal efficiency. We ranked P blastomere-enriched mRNAs based on ribosome occupancy, and identified 19 that ranked in the lowest subset (ribosome occupancy <0.1; Fig. S5). ∼65% of these were among the “P granule transcripts” set defined above, including *cbd-1* which strongly localizes to P granules (Fig. 1, Fig. S2). Among the remainder, three transcripts localized weakly to P granules and exhibited a low ranking in the MEG-3::GFP iCLIPs (*gly-20, ZC155.4* and *R04D3.3,* Fig. S2). We conclude that low ribosome occupancy is one, but likely not the only, criterion for enrichment in P granules.

**Figure 2.**
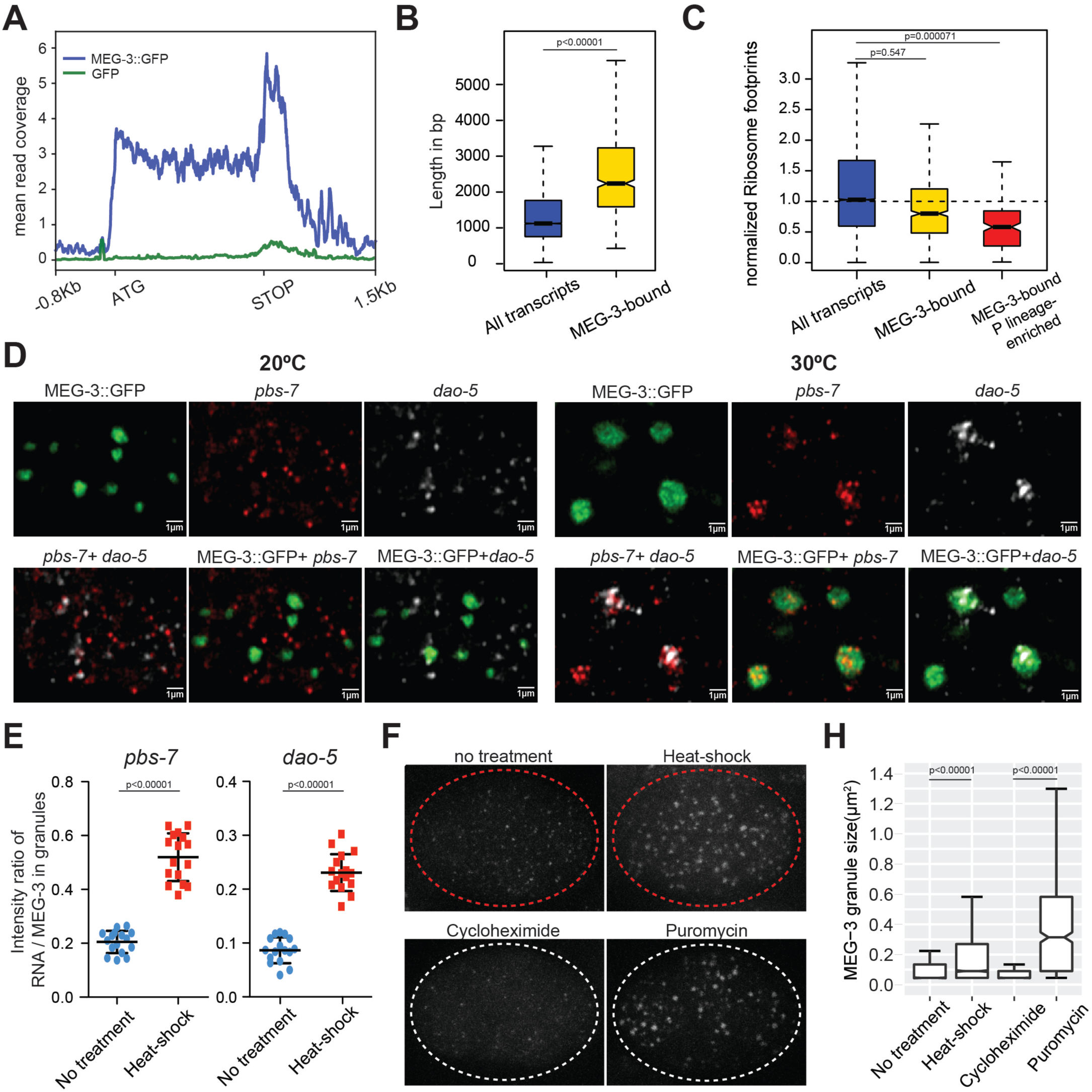
MEG proteins recruit mRNAs into P granules by a sequence non-specific mechanism that favors ribosome-poor mRNAs. (A) Metagene analysis of the distribution of MEG-3::GFP iCLIP reads on the 657 MEG-3-bound transcripts (blue line) compared to GFP iCLIP (green line) reads. (B) Box plot showing the distribution of transcript length for embryonic transcripts (15,345 loci) versus MEG-3-bound transcripts (657 loci). P values were calculated using an unpaired t-test. Each box extends from the 25th to the 75th percentile, with the median indicated by the horizontal line; whiskers extend to the highest and lowest observations. (C) Box plot showing ribosome occupancy in wild-type embryos for three gene categories: embryonic transcripts (15,345 loci), MEG-3-bound transcripts (657 loci) and MEG-3-bound transcripts enriched in P lineage (187 loci). Because ribosome profiling was performed on whole embryos, footprint counts are averages across all cells. Ribosome profiles for the subset of MEG-3::GFP-bound mRNAs that are also enriched in the P lineage (Lee et al., 2017), therefore, are more likely representative of profiles of mRNAs in P granules. See Fig. S4C for ribosome occupancy of P granule transcripts (MEG-3-bound RNAs above the *nos-2* cluster). P values were calculated using an unpaired t-test. See Fig. 2B for box plot description. (D-E) Photomicrographs of P_2_ blastomeres expressing MEG-3::GFP and hybridized to probes against two non-P granule transcripts (*pbs-7* and *dao-5*) before (20°C) and after 15 minutes of heat-shock (30°C). Images are single Z sections and are representative of data quantified in (E). See Fig. S4E for whole embryo images. (E) Graphs showing the intensity ratio of RNA over GFP in MEG-3::GFP granules under no heat-shock (blue dots) or heat-shock (red dots) conditions. Each data point represents the average value for all MEG-3::GFP granules in a single Z section (16 sections were collected from two embryos per condition). P values were calculated using an unpaired t-test. The center horizontal lines indicate the mean and bars represent the SD. (F) Photomicrographs of 4-cell embryos expressing MEG-3::GFP under the indicated treatments. Embryos were derived from *mex-5(RNAi)mex-6(RNAi)* hermaphrodites, which do not localize P granules (Smith et al., 2016). Embryos treated with cycloheximide or puromycin were derived from hermaphrodites also treated with *ptr-2*(RNAi) to permeabilize the egg shells of embryos. Images are representative of data quantified in (H). (H) Box plot showing the size distribution of MEG-3::GFP granules under different treatments as described in F. P values were calculated using an unpaired t-test. Each box extends from the 25th to the 75th percentile, with the median indicated by the horizontal line; whiskers extend to the highest and lowest observations.

### Correlation between P granule exit and translational activation for two mRNAs translated in the germline founder cell P_4_

Among the 18 P granule transcripts analyzed by *in situ* hybridization, we noticed two (*nos-2* and *Y51F10.2*) that transitioned to a more diffuse cytoplasmic localization in the last P blastomere, the germline founder cell P_4_ (Fig. 3A-B). All other transcripts, in contrast, remained in P granules and their levels diminished starting in the P_4_ stage as is typical for maternal mRNAs (Seydoux and Fire, 1994) (Fig. S6A). As mentioned above, *nos-2* codes for a Nanos homolog that becomes translated in P_4_ (Subramaniam and Seydoux, 1999). *Y51F10.2* is a new P granule transcript that had not been characterized before. We tagged the *Y51F10.2* open reading frame with a small epitope by genome editing, and found that like *nos-2*, *Y51F10.2* is translated specifically in P_4_ (Fig. 3A). *nos-2* translation is repressed prior to the P_4_ stage by *mex-3* and activated in P_4_ by *pos-1* (Jadhav et al., 2008). We found that the same is true for *Y51F10.2* (Fig. 3A-B). In *mex-3(RNAi)* embryos, *nos-2* and *Y51F10.2* mRNAs were translated precociously and did not localize to P granules. Conversely, in *pos-1(RNAi)* embryos, *nos-2* and *Y51F10.2* were not translated and remained in P granules in P_4_ (Fig. 3A-B). These observations confirm a link between P granule localization and translational repression. We found, however, that in *meg-3 meg-4* mutants, *nos-2* and *Y51F10.2* translational regulation was not affected despite the absence of granules. *nos-2* and *Y51F10.2* were translationally silent prior to the P_4_ stage and began translation in P_4_ in *meg-3 meg-4* mutants as in wild-type (Fig. 3A-B). Furthermore, MEG-3-bound transcripts as a class maintained low ribosome occupancy in *meg-3 meg-4* mutants as in wild-type (Fig. S6B). These results indicate that neither localization to P granules nor binding to MEG-3 is a requirement for translational silencing or activation. Translation silencing, however, appears to be a requirement for localization to P granules, with translational activation correlating with P granule exit.

**Figure 3.**
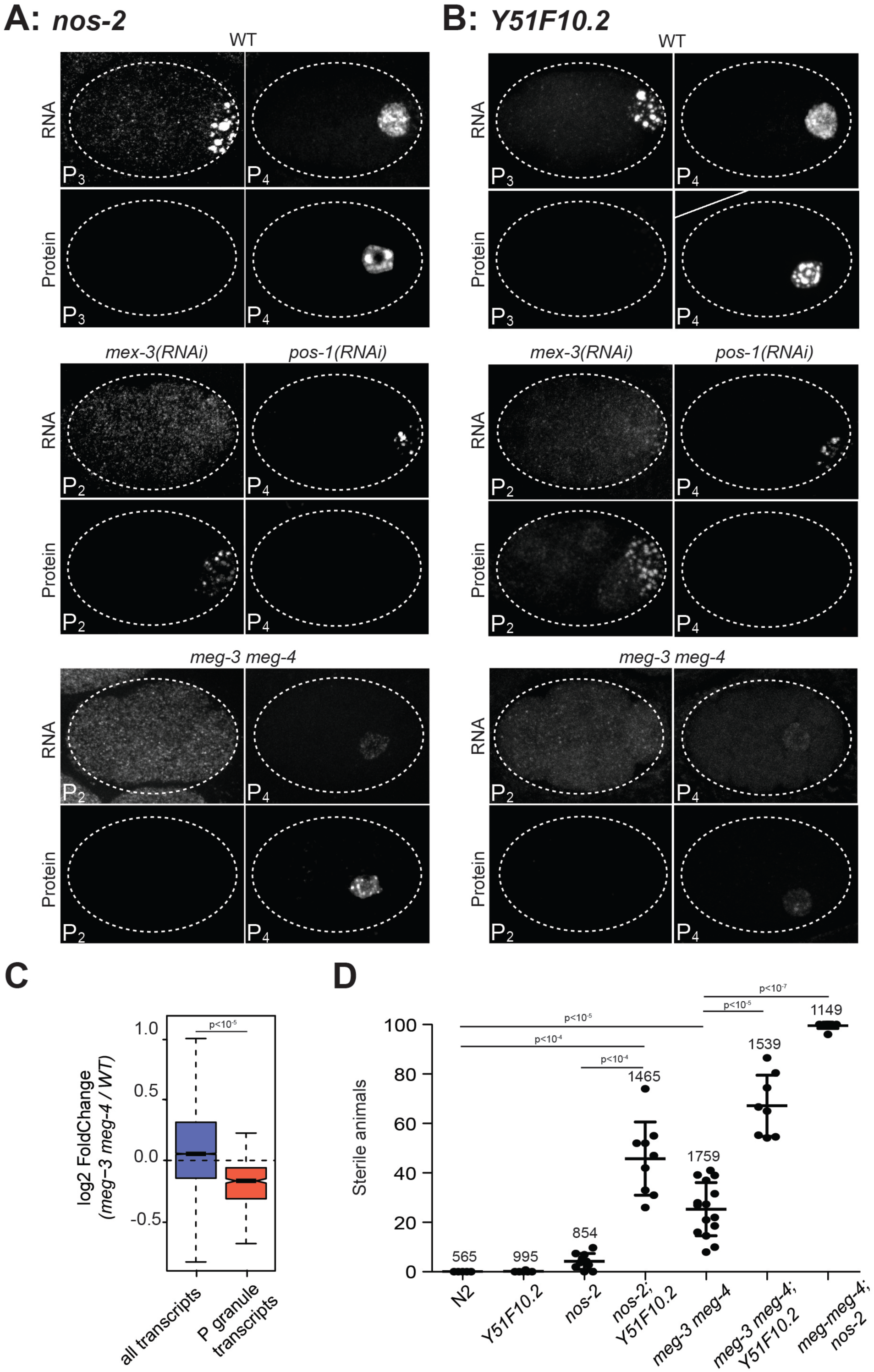
P granules enrich maternal mRNAs required for germ cell development in the nascent germline. (A and B) Photomicrographs of embryos of indicated stages and genotypes and hybridized to fluorescent probes or antibodies to visualize *nos-2* and *Y51F10.2* transcripts and proteins. Embryos expressing NOS-2::3xFLAG and Y51F10.2::OLLAS were used for these experiments. Note the correlation between RNA in granules and no protein expression, and RNA in the cytoplasm and protein expression in wild-type, *mex-3(RNAi)* and *pos-1(RNAi)* embryos. In *meg-3 meg-4* embryos, RNAs are not in P granules and thus are not preferentially segregated to P_4_, resulting in lower RNA levels in that cell. (C) Box plot showing the fold change in expression between wild-type and *meg-3 meg-4* for 15,345 embryonic transcripts and the 492 P granule transcripts. P values were calculated using an unpaired t-test. See Fig. 2B for box plot description. (D) Graphs showing the percentage of sterile animals (no germline) among progeny of hermaphrodites with the listed genotypes (maternal-effect sterility). Each dot represents the brood from a single hermaphrodite allowed to lay eggs for 24 hours. Total number of progeny scored across all broods is indicated for each genotype. P values were calculated using an unpaired t-test. The center horizontal lines indicate the mean and bars represent the SD.

### Role for P granules in enriching mRNAs coding for regulators of germ cell fate

We noticed that *nos-2* and *Y51F10.2* transcript and protein levels were lower in *meg-3 meg-4* mutants compared to wild-type (Fig. 3A-B). At each asymmetric division that gives rise to germline and somatic blastomeres, P granules are segregated preferentially to the germline blastomere (Strome and Wood, 1982) (Fig. S1A). Enrichment in P granules, therefore, is predicted to enrich mRNAs in P_4_. Maternal mRNAs are degraded more rapidly in somatic blastomeres than in germ (P) blastomeres (Seydoux and Fire, 1994), so enrichment in P granules is also predicted to protect mRNAs indirectly from degradation. As predicted, we found that P granule transcripts on average were present at lower levels in *meg-3 meg-4* embryos compared to wild-type as determined by RNAseq (Fig. 3C). 30% of *meg-3 meg-4* mutants develop into sterile adults (Fig. 3D) (Wang et al., 2014), raising the possibility that the lower levels of P granule transcripts in P_4_ compromise germline development. To explore this hypothesis, we examined the effect of combining deletions in *nos-2* and *Y51F10.2* with deletions in *meg-3* and *meg-4*. Embryos derived from mothers carrying deletions in *nos-2* or *Y51F10.2* single mutants were close to 100% fertile (Fig. 3D). In contrast, *nos-2; Y51F10.2* double mutant mothers laid 46±15% sterile progeny that lacked a germline (maternal-effect sterility), suggesting that these maternal products contribute redundantly to germ cell fate specification (Fig. 3D). Remarkably, the majority of *nos-2*; *meg-3 meg-4* and *Y51F10.2*; *meg-3 meg-4* triple mutants grew into sterile animals lacking a germline (Fig. 3D). This synthetic effect is consistent with a role for *meg-3* and *meg-4* in concentrating transcripts coding for germ cell fate regulators, including *nos-2*, *Y51F10.2* and likely others, beyond a threshold required for robust germline development. This analysis also identifies Y51F10.2 as a new maternal factor promoting germ cell fate. Y51F10.2 is the *C. elegans* orthologue of human TRIM32, a member of the broadly conserved TRIM-NHL protein family implicated in protein ubiquitination and mRNA regulation (Tocchini and Ciosk, 2015). It appears unlikely, however, that all P granule transcripts function in germ cell fate specification, as many turn over before ever releasing from P granules (Fig. S6), and at least some have known functions in unrelated processes (e.g. *cbd-1;* a chitin-binding protein required for egg shell biogenesis (Johnston et al., 2010)).

### Condensation of RNA with MEG-3 into nanoscale gels

Our observations indicate that mRNAs are recruited into P granules in a sequence non-specific manner as part of the MEG phase. We showed previously that recombinant MEG-3 readily phase separates *in vitro* with total *C. elegans* RNA (Putnam et al., 2019). To investigate whether MEG-3 shows any bias when presented with specific sequences, we synthesized 9 fluorescently labeled RNAs (800-1300nt size range) corresponding to embryonic transcripts with strong, intermediate, or no localization to P granules *in vivo* under normal culture conditions. Each transcript was combined with recombinant His-tagged MEG-3 in condensation buffer containing 150mM salt. In the absence of RNA, MEG-3 formed irregular aggregates with a broad size range (Fig. 4A,B). Addition of RNA led to the formation of uniform MEG-3/RNA condensates with radii of less than 400 nm in size (Fig. 4A,B and Fig. S7A,B), consistent with the size of MEG-3 condensates *in vivo* (Putnam et al., 2019). We found that all 9 transcripts stimulated MEG-3 condensation and became enriched in the condensates with similar efficiencies (Fig. 4C, Fig. S7C).

**Figure 4.**
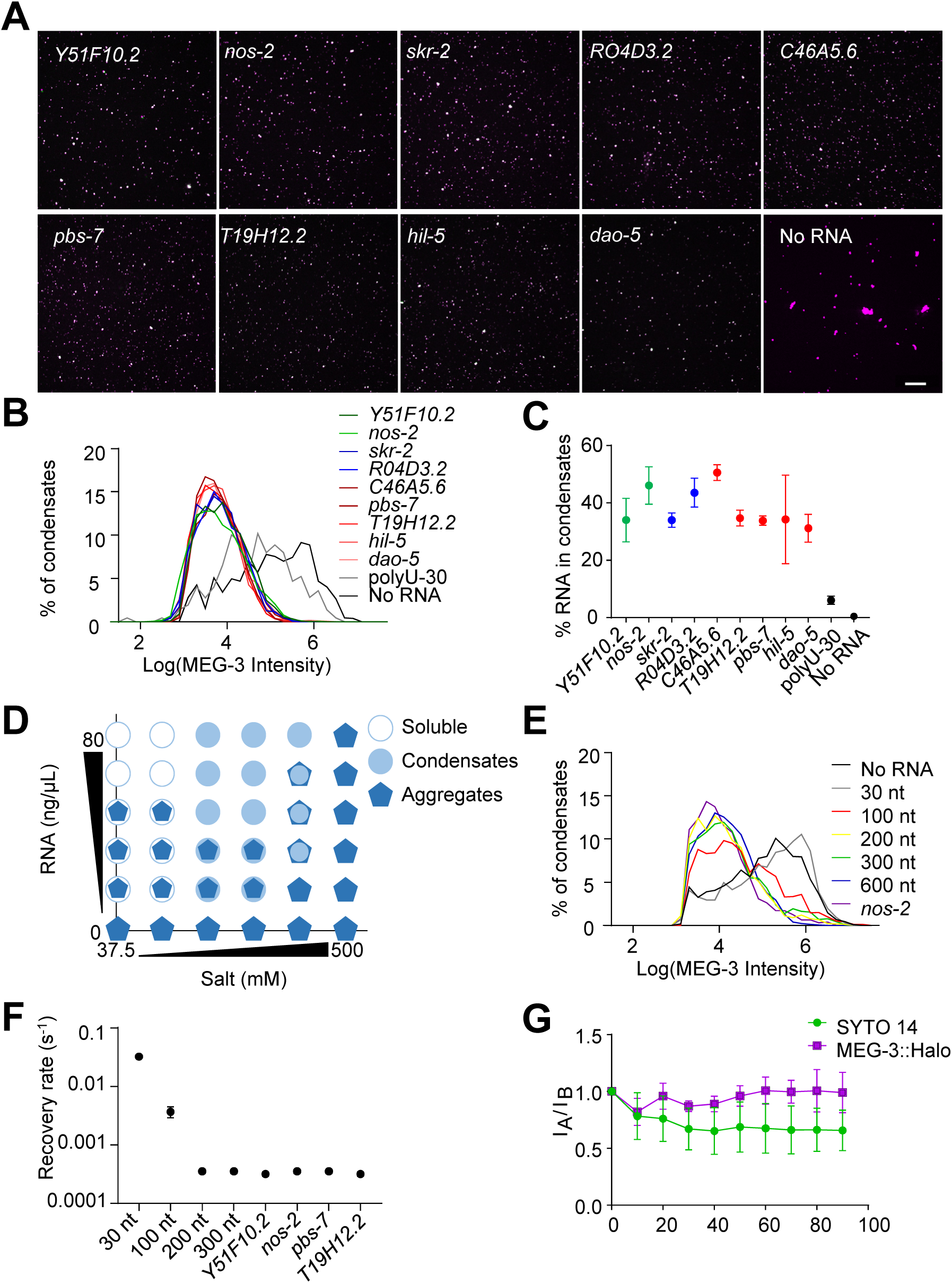
Long RNAs stably associate with the MEG gel phase. (A) Representative photomicrographs of condensates of MEG-3 and indicated RNA after incubation in condensation buffer. Reactions contained 500 nM MEG-3 and 20 ng/µL *in vitro* transcribed RNA. MEG-3 (magenta) was trace labeled with Alexa647 and RNA (green) was trace labeled with Alexa546 (Methods). Scale bar is 20 µm. (B) Histograms of MEG-3 intensity (log10 scale) normalized to the total number of condensates in each reaction assembled as in A for each RNA indicated. Each histogram includes condensates from 12 images collected from 3 experimental replicates. RNAs correspond to transcripts with MEG-3 iCLIP counts above the *nos-2* cluster (*nos-2, Y51F10.2*), below the *nos-2* cluster (*skr-2, R04D3.2*) and not recovered in the MEG-3 iCLIPs (*C46A5.6, pbs-7, T19H12.2, hil-5, dao-5*). (C) Graph showing the percent of RNA fluorescence in MEG-3 condensates compared to total RNA assembled as in A. Each data point represents condensates from 12 images collected from 3 experimental replicates. Circles indicate the mean and bars represent the SD. RNAs corresponding to transcripts with MEG-3 iCLIP counts above the *nos-2* cluster (green), below the *nos-2* cluster (blue) and not recovered in the MEG-3 iCLIP (red). (D) Phase diagram of MEG-3 condensate composition under varying RNA and salt concentrations. For representative images and quantitation corresponding to positions in the diagram refer to Fig. S7D-G. MEG-3 was present in three states: i) soluble MEG-3 (no condensates detected, open circles), ii) small uniform condensates (Log(I)≤4.6, filled circles), iii) large irregular condensates (Log(I)>4.6, pentagons). In conditions with mixed MEG-3 states, the larger object represents the predominant population. (E) Histograms of MEG-3 intensity (log10 scale) normalized to the total number of condensates/aggregates in each reaction assembled as in Fig. S8A. Each histogram includes data from 12 images collected from 3 experimental replicates. Lines represent reactions assembled for the indicated RNA. (F) Graph showing rates of fluorescence recovery after photobleaching (FRAP) for indicated RNAs in MEG-3 condensates. Values were normalized to initial fluorescence intensity, corrected for photobleaching and plotted. Circles indicate the mean (n=6) and bars represent the SD. Refer to Fig. S8C,D for representative images and time traces. (G) Graphs showing the fraction of MEG-3::Halo or SYTO 14 retained in the condensate phase after extrusion from embryos normalized to the fraction before extrusion (time 0). Total Halo or SYTO 14 fluorescence in granules was measured before laser puncture (I_B_) and after laser puncture (I_A_), corrected for photobleaching and used to calculate a fluorescence ratio (I_A_/I_B_). Means are indicated along with error bars representing ± SD calculated from 5 embryos. Refer to Fig. S9A,B for representative images.

MEG-3 has a predicted pI of 9.3 (ExPASy) and thus could potentially interact with the sugar-phosphate backbone of RNA through electrostatic interactions. To test this hypothesis, we examined MEG-3 condensation behavior in the presence of varying concentrations of salt and RNA (*nos-2* transcript). In this analysis, we distinguished aggregates from condensates based on size as determined in Fig. 4B. In the absence of RNA, MEG-3 formed aggregates under all salt concentrations tested. In high salt (500mM) conditions, MEG-3 continued to form aggregates even in the presence of *nos-2* RNA and these aggregates did not recruit *nos-2* RNA. Decreasing salt concentrations and increasing RNA concentrations shifted the balance from MEG-3 aggregates to MEG-3/RNA condensates. Remarkably, at the lowest concentrations of NaCl, high concentrations of RNA caused MEG-3 to solubilize with no visible aggregates or condensates (Fig. 4D and Fig. S7D-G). These observations are consistent with MEG-3 interacting with RNA in a salt-sensitive manner and suggest that electrostatic interactions with RNA compete with the MEG-3/MEG-3 interactions that lead to aggregation.

To determine whether RNA length affects MEG-3 condensation behavior, we tested RNAs of varying sizes in the condensation assay. We found that short RNAs (30 and 100 nt) were not as efficient as longer RNAs at stimulating MEG-3 condensation (Fig. 4E, Fig. S8A,B). We previously showed that MEG-3 protein becomes immobilized in MEG-3/RNA condensates, with no MEG-3 exchange detected within one minute of condensation (Putnam et al., 2019). To determine whether RNAs also become trapped in MEG-3 condensates, we performed FRAP experiments to measure the rate of RNA exchange between dilute and condensed phases comparing RNAs of different lengths. We found that short RNAs (30 and 100 nt) were mobile in MEG-3 condensates. In contrast, longer RNAs (200nt and higher), including 4 full-length transcripts, exhibited no detectable exchange (Fig 4F, Fig. S8C,D). We conclude that MEG-3 interacts most efficiently with mRNA-sized RNAs, which become trapped in the MEG-3 condensates.

To examine whether mRNAs also associate stably with the MEG phase of P granules *in vivo*, we labeled permeabilized live embryos with the RNA dye SYTO 14. As expected, we observed intense SYTO 14 fluorescence in MEG-3-positive granules (Fig. S9A,B). We verified *in vitro* that SYTO 14 fluorescence is sensitive to RNA and does not interact significantly with MEG-3 or PGL-3 in the absence of RNA (Fig. S9C). When released from embryos by laser puncture of the eggshell, MEG-3 granules remain stable in aqueous buffer, whereas PGL-1 and PGL-3 dissolve immediately (Putnam et al., 2019). We found that MEG-3 granules remained positive for SYTO 14 *ex vivo* for over 1.5 minutes (the maximum time tested; Fig. 4G, Fig. S9B). These observations indicate that mRNAs associate stably with the MEG phase *ex vivo* as observed *in vitro*, and confirm that the majority of RNAs in embryonic P granules do not reside in the PGL phase.

### RNA-dependent clustering of MEG-3 condensates into super-assemblies

When assembled in the presence of PGL-3, MEG-3/RNA condensates recruit PGL-3 and accumulate on the surface of PGL-3 droplets forming a discontinuous porous shell (Putnam et al., 2019). This architecture is also observed *in vivo* during P granule assembly in early embryos (Putnam et al., 2019). *In vitro* and *ex vivo*, treatment of the MEG-PGL co-condensates with 1M NaCl dissolves the central PGL core, leaving behind a hollow shell of coalesced MEG-3/RNA condensates (Putnam et al., 2019). Using the *nos-2* transcript, we confirmed that RNA remains trapped in the MEG-3 phase after salt treatment and that MEG-3/*nos-2* RNA condensates form stable super-assemblies that persist even in the absence of the PGL phase (Fig. 5A, Fig. S10). MEG-3/poly-U30 condensates also assembled on the surface of PGL condensates but behaved differently upon salt treatment: poly-U30 was removed from the MEG-3 condensates and the MEG condensates dissociated into individual condensates that did not maintain the hollow shell configuration observed with MEG-3/*nos-2* RNA condensates (Fig. 5B). This observation suggested that RNA modulates the ability of MEG condensates to form super-assemblies. To test this hypothesis directly, we assembled the MEG-PGL co-condensates without RNA. We found that, in the presence of PGL-3, formation of large MEG-3 aggregates is inhibited and MEG-3 forms small condensates on the surface of the PGL-3 droplets in a similar configuration to that of co-condensates formed with RNA, except that PGL-3 becomes even more enriched in the MEG-3 condensates (Fig. 5C). Salt treatment of the RNA-free co-condensates induced dissolution of the PGL phase and, interestingly, caused the MEG condensates to disperse into individual condensates, as observed with poly-U (Fig. 5C). We conclude that long RNA molecules that interact stably with the MEG condensates are required for MEG condensates to link to each other to form super-assemblies.

**Figure 5.**
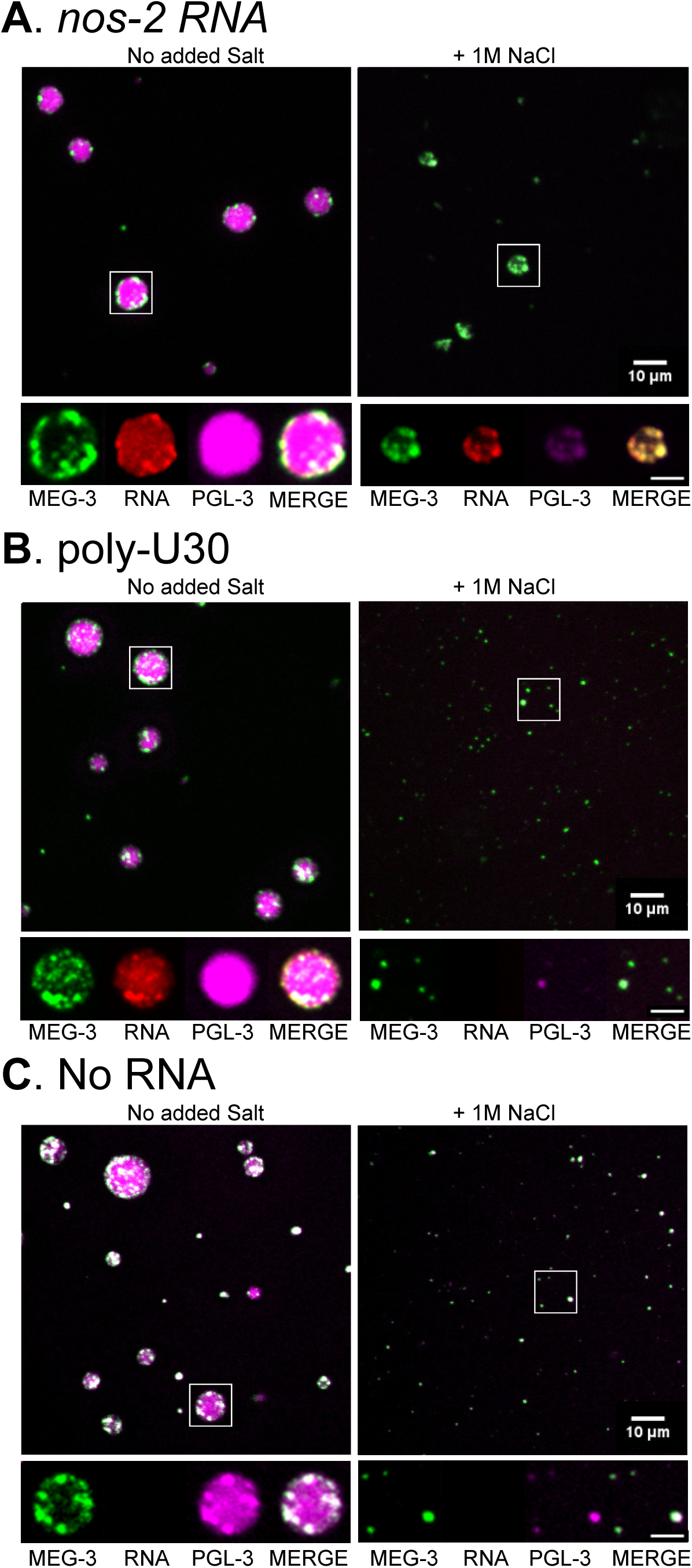
RNA promotes the formation of super-assemblies of MEG condensates. (A) Top panel: Representative photomicrograph of condensates of MEG-3, PGL-3 and *nos-2* RNA incubated for 30 min in condensation buffer. NaCl (final 1M) was added as indicated for an additional 15 min. Reactions contained 500 nM MEG-3, 5 µM PGL-3 and 20 ng/µL *nos-2* RNA. MEG-3 (green) was trace labeled with Alexa555, PGL-3 (magenta) was trace labeled with Alexa647, *nos-2* RNA (red) was trace labeled with Alexa488. Scale bar is 10 µm. Bottom panel: high magnification photomicrographs of the highlighted condensates in main panel showing MEG-3, PGL-3, RNA and merge views. Scale bar is 5 µm. Refer to Fig. S10 for additional images. (B) Same as in A but condensates were assembled with poly-U30 RNA. (C) Same as in A but condensates were assembled in the absence of RNA. Note that PGL-3 is recruited to the MEG phase under all conditions, but most strongly in the absence of RNA.

### Parallels between P granules and stress granules and a revised model for the assembly of cytoplasmic RNA granules

In this study, we report on the transcriptome and assembly mechanism of P granules, a classic model for cytoplasmic RNA granules. Our findings reveal parallels between P granules and stress granules, RNA granules that assemble under conditions of energy stress. As reported for stress granules (Aulas et al., 2017; Kedersha et al., 2000; Khong et al., 2017), we find that P granule assembly favors long mRNAs that are ribosome-depleted and is enhanced by treatments that release mRNAs from polysomes (heat-shock and puromycin). For two mRNAs coding for germ cell fate regulators, we show that translational repression is a requirement for P granule entry, and translational activation correlates with P granule exit. Recruitment of mRNAs to stress granules is an inefficient process, with only a minority of molecules for most mRNA species localizing in the granules (Khong et al., 2017). Recruitment into P granules is also inefficient with only ∼30% of molecules localizing to granules for two of the most robust P granule transcripts. Finally, P granules contain proteins also found in stress granules (poly-A binding protein, TIA-1 and the DDX3/LAF-1 RNA helicase) and interact with P bodies, as do stress granules (Elbaum-Garfinkle et al., 2015; Gallo et al., 2008; Shih et al., 2012). We suggest that embryonic P granules are a developmentally-regulated version of stress granules. Stress granules have been proposed to serve as temporary storage hubs for mRNAs awaiting translational activation post-stress (Buchan and Parker, 2009; Ivanov et al., 2019). Our findings suggest that P granules may perform a similar function for at least of subset of P granule mRNAs coding for germ cell fate regulators that are translated in the germline founder cell P_4_. Because of their spatially regulated assembly (Putnam et al., 2019; Smith et al., 2016), P granules also promote the preferential segregation of RNAs coding for germ cell regulators to the germline founder cell, thus ensuring robust germ cell fate specification.

Stress granule assembly has been proposed to depend on low affinity RNA:protein and RNA:RNA interactions that collectively drive liquid-liquid phase separation of mRNA molecules with RNA-binding proteins (Ditlev et al., 2018; Protter and Parker, 2016; Van Treeck et al., 2018; Van Treeck and Parker, 2018). Our findings, however, demonstrate that mRNAs are not recruited as part of the liquid phase of P granules. Instead, mRNAs are recruited by gelation with the intrinsically-disordered protein MEG-3 (and its paralog MEG-4). MEG-3 condenses in a sequence non-specific manner with RNA to form nanoscale gels. The gels associate with the surface of the PGL-rich liquid core of P granules to form super-assemblies that link via RNA-dependent interactions. Super-resolution microscopy has revealed that stress granules are also super-assemblies of smaller condensates that reversibly cluster during granule assembly and disassembly (Cirillo et al., 2019; Niewidok et al., 2018; Wheeler et al., 2016). Furthermore single-molecule analyses have shown that mRNAs are non-dynamic when inside stress granules (Moon et al., 2018). These and our findings suggest a revised model for RNA granule assembly. mRNAs are recruited into RNA granules by sequence non-specific gelation with intrinsically-disordered proteins to yield nanoscale gel condensates. The gel condensates favor phase separation of RNA-binding proteins into “client” liquid phases, such as the PGL liquid core of P granules (Putnam et al., 2019) and the liquid shell of stress granules (Protter and Parker, 2016). The client liquid phases, in turn, contribute to RNA granule assembly by providing surfaces onto which nanoscale RNA gels cluster to form super-assemblies. In this revised model, RNA granule disassembly involves both dispersal of super-assemblies into individual nanoscale gel condensates and dissolution of liquid phases. Identification of the mechanisms that regulate the clustering and dispersal dynamics of nanoscale RNA gels will be key to understanding the mesoscale liquid-like behavior of RNA granules.

## Acknowledgments

We thank the Johns Hopkins Neuroscience Research Multiphoton Imaging Core (NS050274) and the Johns Hopkins Integrated Imaging Center (S10OD023548) for excellent microscopy support. We thank the Lavis lab for Halo Ligand JF_646_, the Griffin lab for MEG-3::Halo strain, Addgene for TEV protease from the Waugh lab, and Colin Wu for assistance with ribosome profiling experiments. We especially thank Dr. Sijung Yun (Yotta Biomed) for his assistance with data analysis. We also thank the Baltimore Worm club and the Seydoux lab for many helpful discussions. This work was supported by the National Institutes of Health (NIH) (grant number R37 HD37047). G.S. is an investigator of the Howard Hughes Medical Institute. Some strains were provided by the CGC, which is funded by NIH Office of Research Infrastructure Programs (P40 OD010440). Data acquired using the Zeiss LSM 800 Confocal reported in this publication was supported by Office of the Director, NIH (OD) of the National Institutes of Health (award number S10OD016374).

## Author contributions

CYL, AAP, SH, TL, JPO conducted the experiments; CYL, AAP, TL and GS analyzed the data. CYL, AAP and GS designed the experiments and wrote the paper.

## FIGURES

**Figure S1.**
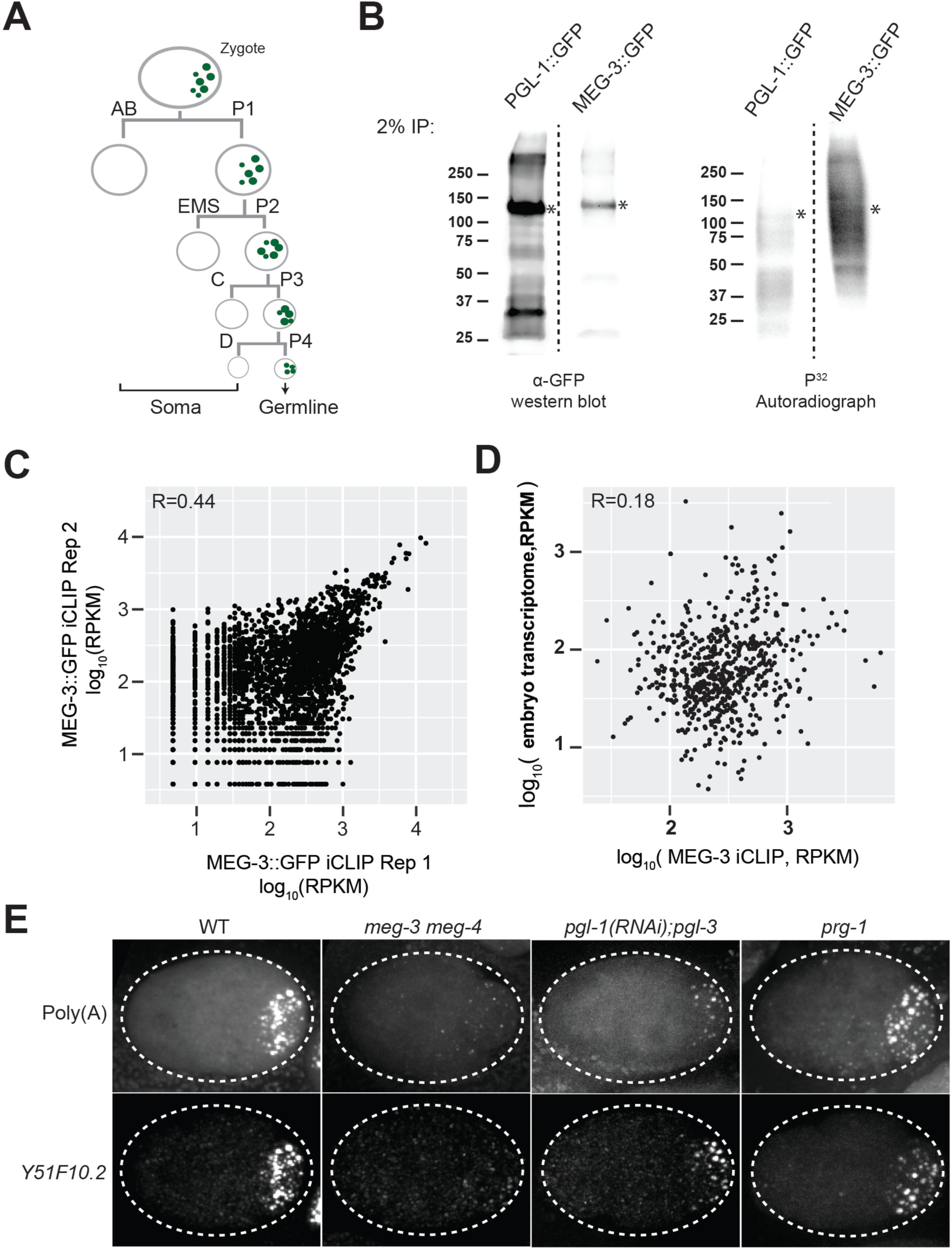
mRNAs are recruited into P granules by the MEG phase. (A) Abbreviated embryonic lineage showing the somatic (AB, EMS, C, and D) and germline (P) blastomeres. Green dots represent P granules. (B) Anti-GFP western blot and autoradiograph of crosslinked anti::GFP immunoprecipitates separated by SDS-PAGE. The immunoprecipitates were 5’ end radiolabeled to visualize cross-linked protein:RNA complexes. * indicates predicted MEG-3::GFP and PGL-1::GFP bands based on respective sizes. Note that PGL-1::GFP protein is more abundant than MEG-3::GFP, yet does not cross-link with RNA as efficiently. C) Graph plotting reads for each transcript (black dots) in MEG-3::GFP iCLIP replicate 1 versus replicate 2. R is the Pearson correlation coefficient. (D) Graph plotting 657 MEG-3-bound transcripts (black dots) with respect to average read count in the MEG-3::GFP iCLIPs (average RPKM of two replicates, X axis) versus transcript abundance in early embryos (Y axis). R is the Pearson correlation coefficient. (E) Photomicrographs of 4-cell embryos of the indicated genotypes hybridized with smFISH probes against poly(A) and *Y51F10.2*. P granule RNA clusters are smaller in *pgl-1(RNAi); pgl-3* embryos, consistent with clustering of MEG-3 condensates to the surface of PGL condensates in wild-type embryos (Putnam et al., 2019) (Fig. 5). The identity of the small polyA clusters in the P_2_ and EMS blastomeres in *meg-3 meg-4* mutant embryos is not known, but may be correlated to the higher levels of maternal RNAs in these cells compared to the anterior somatic blastomeres where many maternal mRNAs are rapidly degraded.

**Figure S2.**
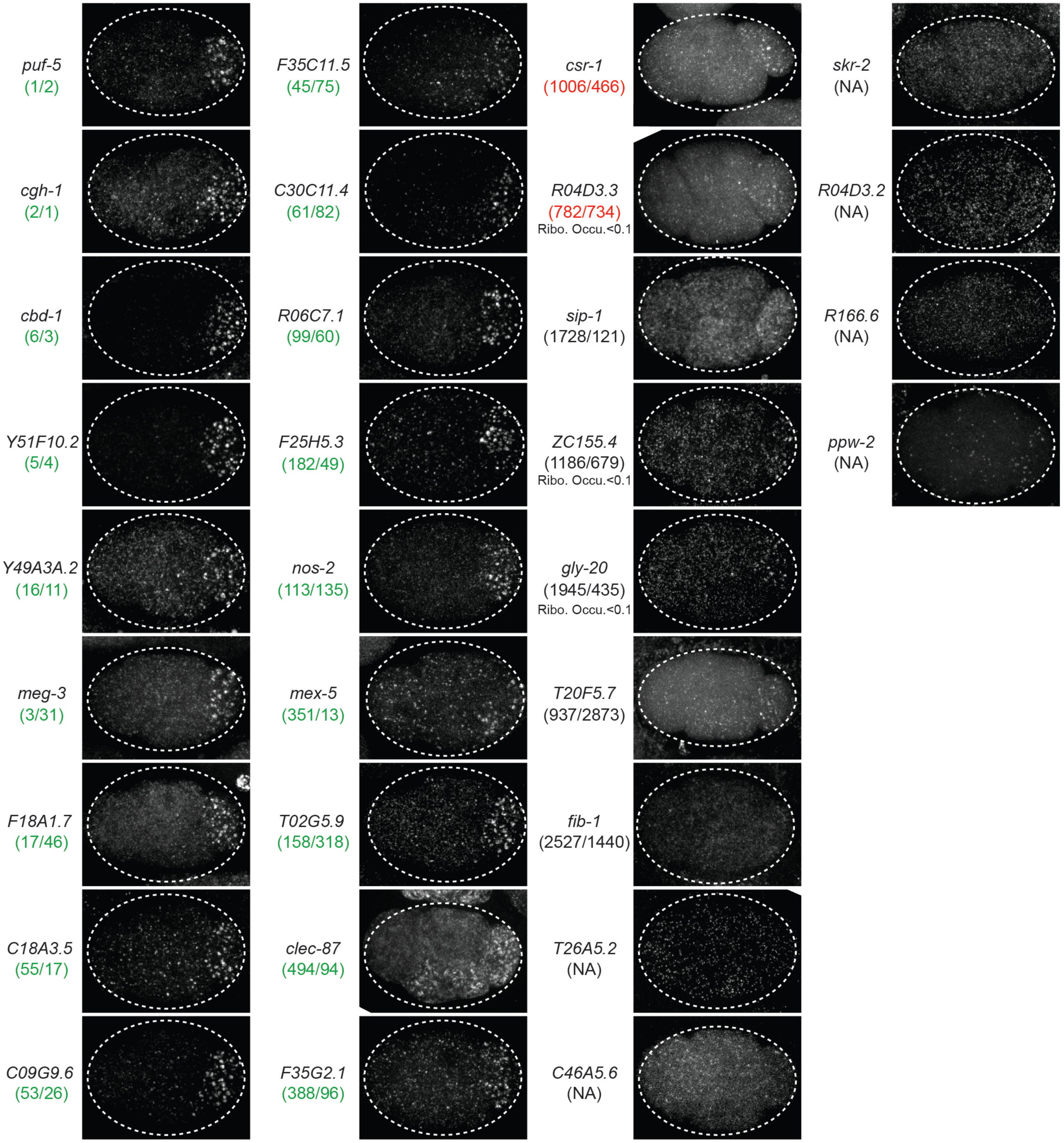
*In situ* hybridization. Photomicrographs of 4-cell wild-type embryos hybridized with smFISH probes as indicated. Maximum Z projections of photomicrographs from at least 10 embryos were examined per probe set. Numbers in parenthesis indicate rank order in two MEG-3::GFP iCLIP experiments. Green indicate rankings at or above the *nos-2* cluster threshold (“P granule transcript” category). Red indicate rankings below the *nos-2* cluster threshold (“MEG-3-bound transcript category”. Black indicate ranking below the GFP background threshold. “NA” indicates no read count in one or both MEG-3::GFP iCLIP experiments.

**Figure S3.**
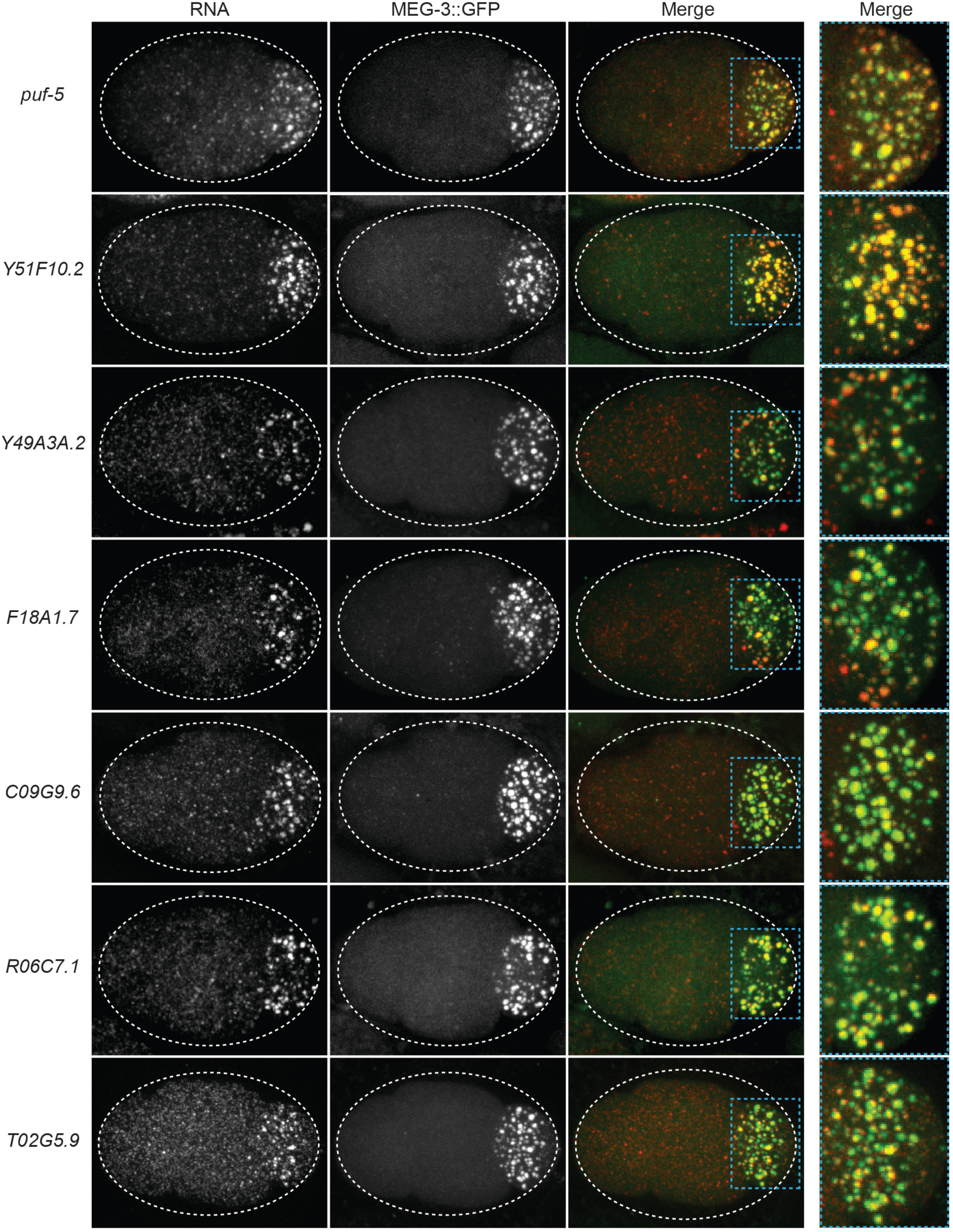
*In situ* hybridization with MEG-3::GFP. Photomicrographs of embryos expressing MEG-3::GFP hybridized with single molecule fluorescence (smFISH) probes as indicated. All 7 are “P granule transcripts” localizing to P granules (MEG-3::GFP), as shown in the higher magnification merged panels.

**Figure S4.**
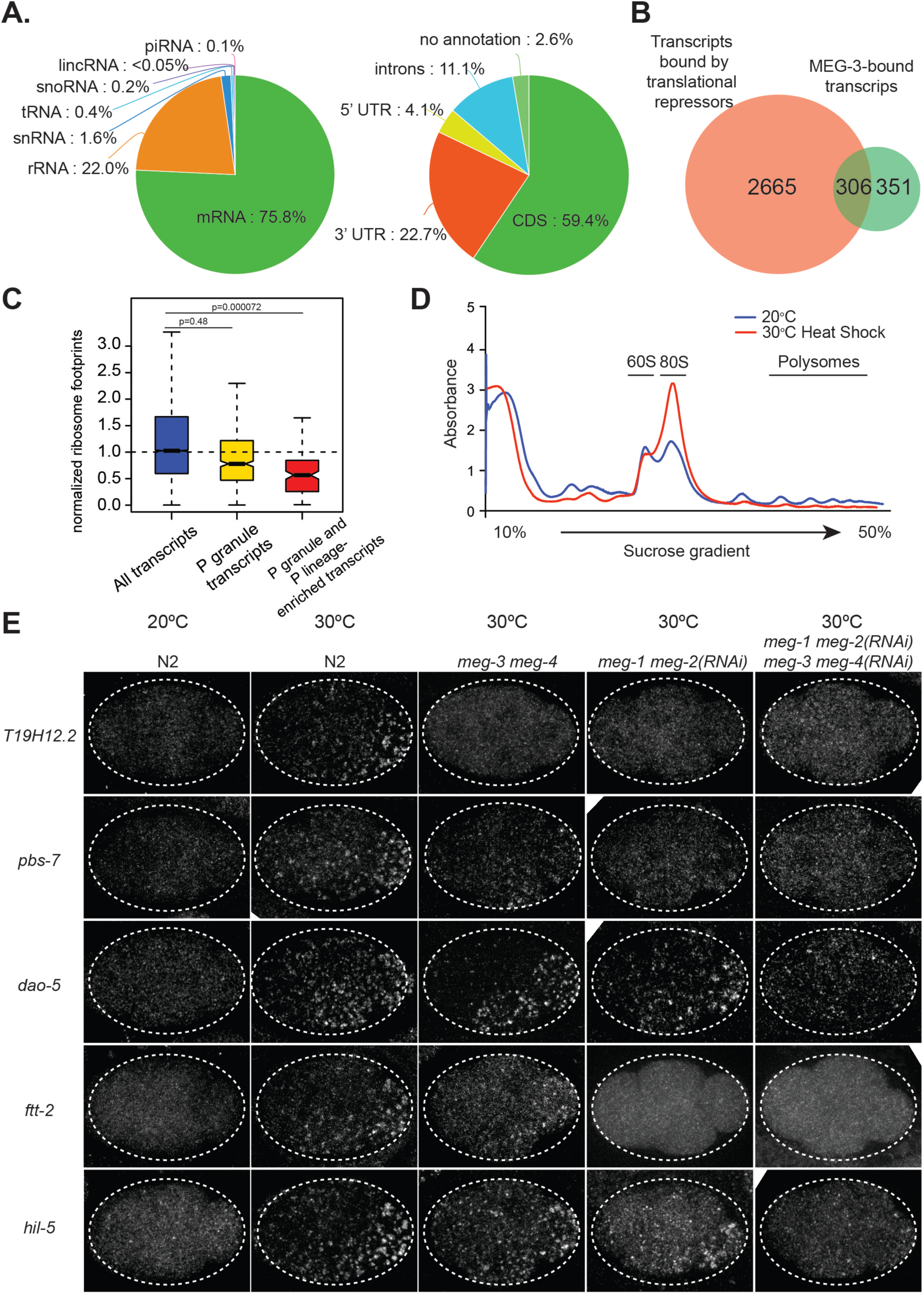
MEG-3 binds ribosome-poor transcripts. (A) First pie chart shows distribution of MEG-3::GFP reads relative to different transcript types (total: 521,189 reads). Second pie chart shows distribution of MEG-3::GFP reads mapping to mRNAs relative to gene features (total: 275,083 reads). (B) Venn diagram showing the overlap between 2,971 transcripts bound by translational repressors (GLD-1, CHG-1, OMA-1 and LIN-4 (Boag et al., 2008; Scheckel et al., 2012; Tsukamoto et al., 2017)) and the 657 MEG-3-bound transcripts defined in this study. Note that the transcripts bound by translation repressors were defined in oocytes whereas the MEG-3-bound transcripts were defined in embryos, therefore 100% overlap is not expected. (C) Same as in Fig. 2C but for the 492 P granule transcript gene set. Because ribosome profiling was performed on whole embryos, footprint counts are averages across all cells. Ribosome profiles for MEG-3-bound, P lineage-enriched transcripts (Lee et al., 2017) are more representative of profiles of mRNAs in P granules bound by MEG-3. P values were calculated using an unpaired t-test. See Fig. 2B for box plot description. (D). Graph showing RNA absorbance of embryonic lysates separated on a sucrose gradient. The lysates were isolated from wild-type embryos raised at 20°C without heat-shock or with heat-shock (30°C for 15 minutes). (E) Photomicrographs of 4-cell stage embryos of indicated genotypes after no heat-shock (20°C) or after heat-shock (30°C) and hybridized to smFISH probes against the indicated transcripts. Maximum Z projection of photomicrographs from at least 10 embryos were examined per probe set. Note that for *dao-5*, *ftt-2* and *hil-5*, RNA granules still form preferentially in P_2_ and EMS in *meg-3 meg-4* mutants. This pattern depends on *meg-1* and *meg-2*, two MEG paralogs (Wang et al., 2014).

**Figure S5.**
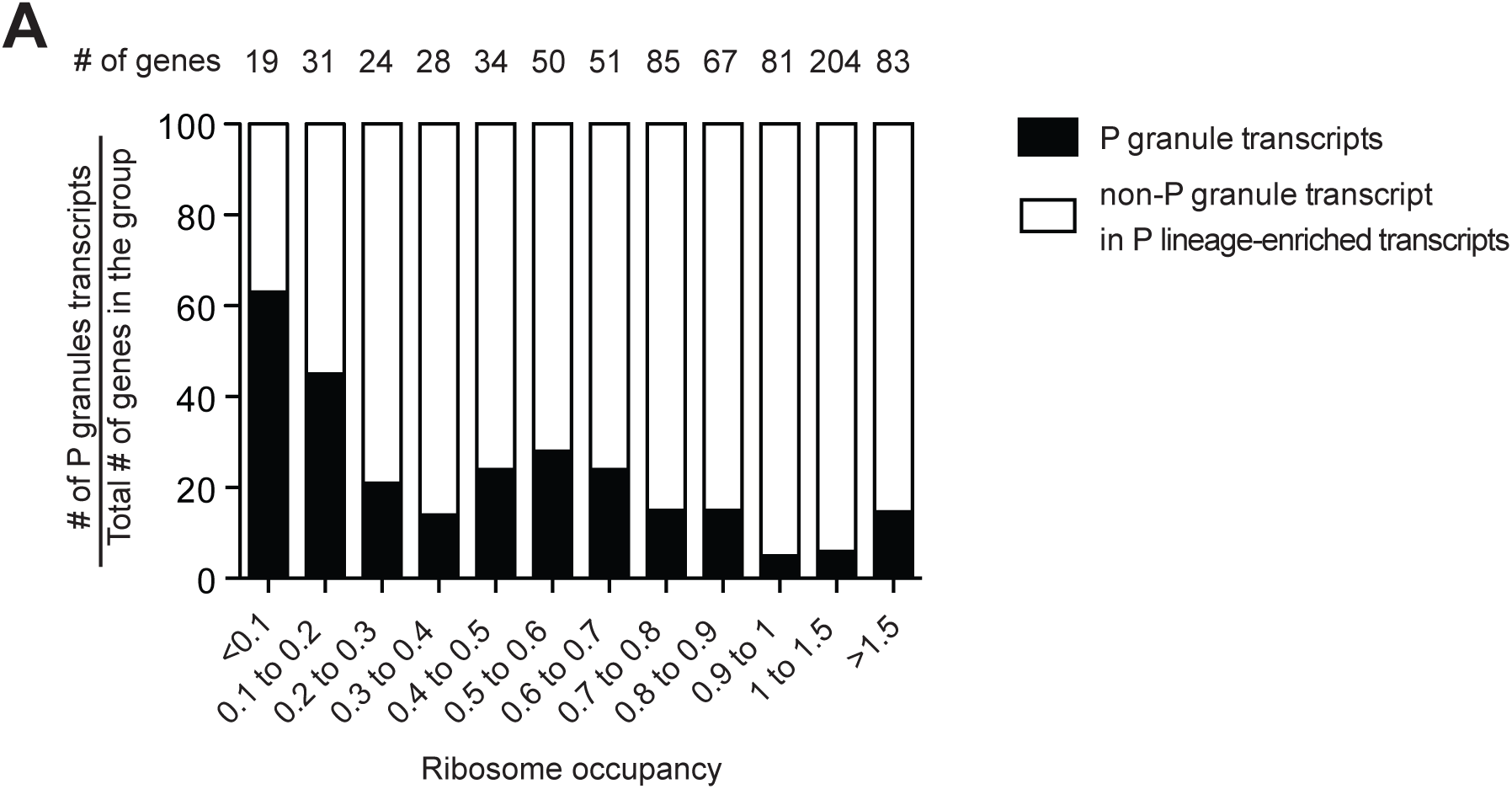
Distribution of P granule transcripts and MEG-3-bound transcripts across ribosome-occupancy classes. Bar graph showing the fraction of P granule transcripts (black) binned by ribosome occupancy. Only P lineage-enriched transcripts (FPM > 25) were included in this analysis (Lee et al., 2017). Total number of loci per bin is indicated.

**Figure S6.**
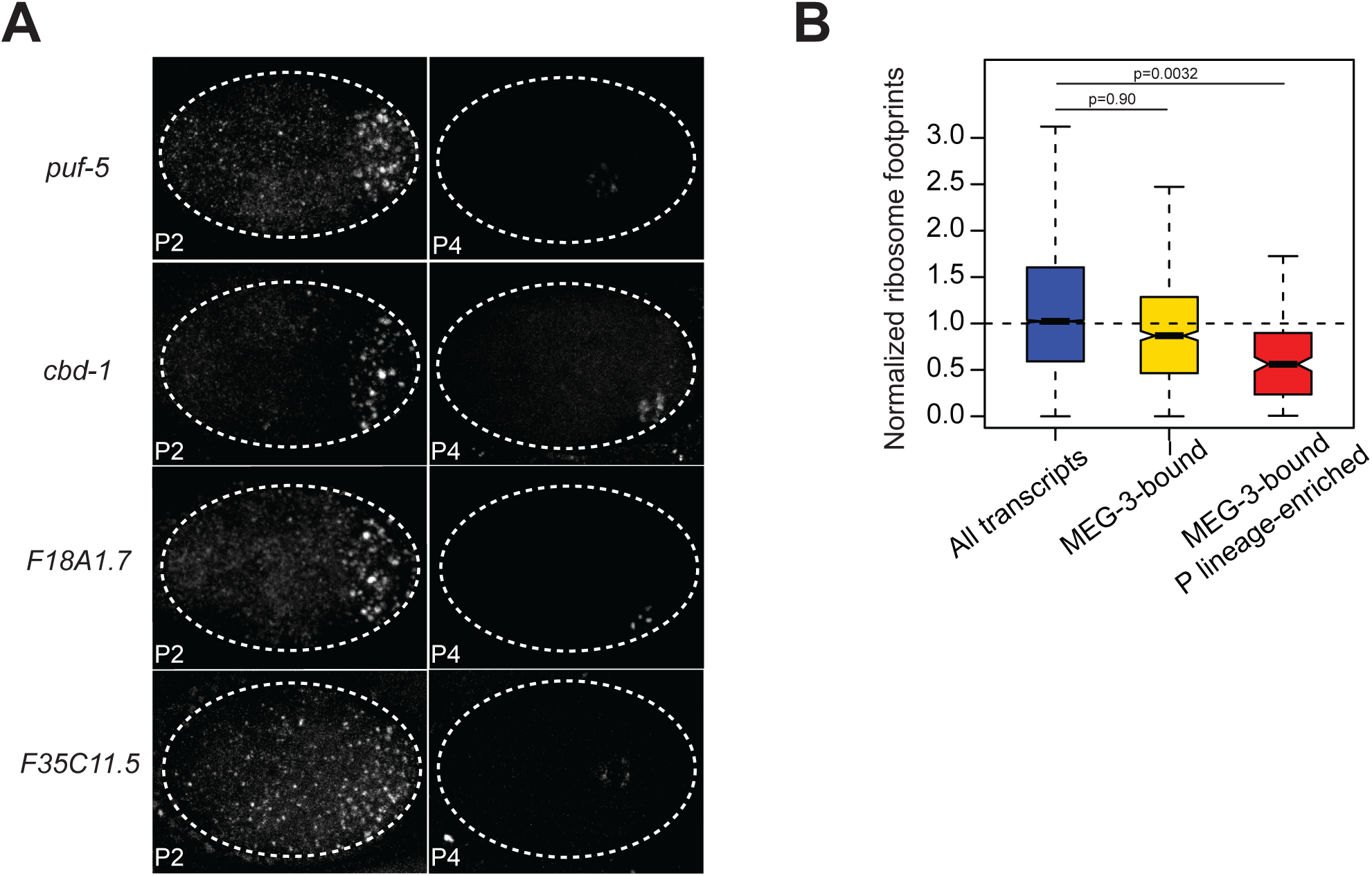
P granule RNA distribution in 4-cell and 28-cell embryos. (A) Photomicrographs of P_2_ and P_4_ stage embryos hybridized with smFISH probes as indicated. Unlike *nos-2* and *Y51F10.2* shown in Figure 3, the transcripts shown here remain associated with P granules through the P_4_ stage and become less abundant overtime. (B) Same as in Fig. 2C but for *meg-3 meg-4* embryos. Because ribosome profiling was performed on whole embryos, footprint counts are averages across all cells. Ribosome profiles for MEG-3-bound, P lineage-enriched transcripts(Lee et al., 2017) are more representative of profiles of mRNAs in P granules bound by MEG-3. P values were calculated using an unpaired t-test. See Fig. 2B for box plot description.

**Figure S7.**
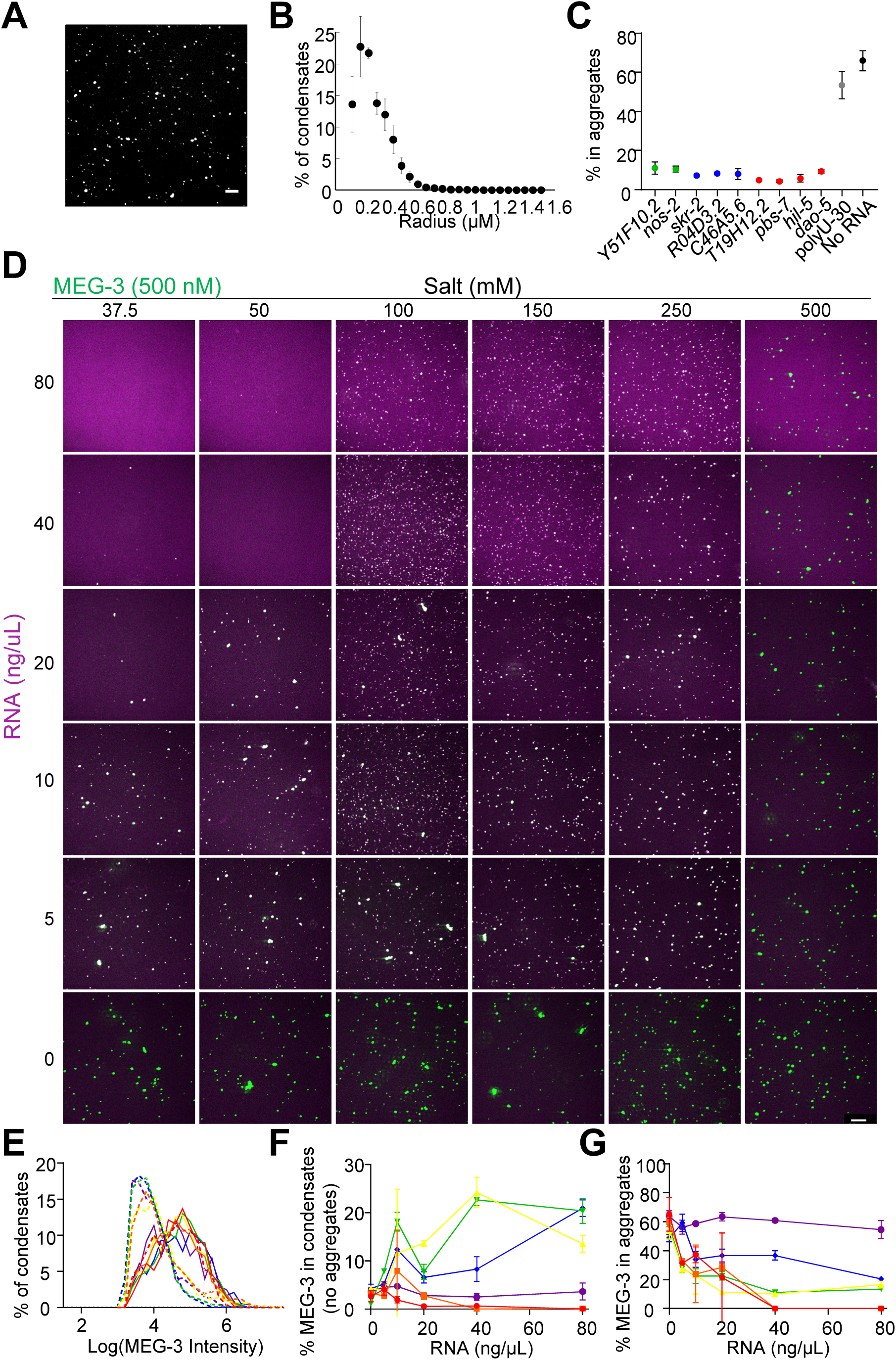
RNA modulates MEG condensation dependent on salt concentration. (A) Representative photomicrograph of condensates of MEG-3 and indicated RNA after incubation in condensation buffer and imaged with a 100X objective. Reactions contained 500 nM MEG-3 and 40 ng/µL *nos-2*. MEG-3 (white) was trace labeled with Alexa647 and RNA (not shown) was trace labeled with Alexa488. Scale bar is 5 µm. (B) Histograms of MEG-3 condensate radii normalized to the total number of condensates in each reaction assembled as in Fig. S7A. Each point represents data from 64 images collected from 4 experimental replicates. Circles indicate the mean and bars represent the SD. (C) Percent of MEG-3 in aggregates for each RNA indicated assembled as in Fig.4A-C. Aggregates were defined as objects with a MEG-3 intensity of Log (I) > 4.6 from histograms in (B) (Methods). Each data point represents data from 12 images collected from 3 experimental replicates. Circles indicate the mean and bars represent the SD. (D) Representative photomicrographs of MEG-3/*nos-2* RNA condensation reactions. Reactions contained 500 nM MEG-3 and indicated concentration of *nos-2* RNA, and salt. MEG-3 was trace labeled with Alexa647 (green) and RNA was trace labeled with Alexa488 (magenta). Scale bar is 20 µm. (E) Histograms of MEG-3 intensity (log10 scale) normalized to the total number of condensates/aggregates in each reaction assembled as (D). Each histogram includes data from 8 images collected from 2 experimental replicates. Lines represent reactions assembled in 150 mM salt (dashed lines) or 500 mM salt (solid lines) for RNA concentrations of 0 ng/µL (red), 5 ng/µL (orange), 10 ng/µL (yellow), 20 ng/µL (green), 40 ng/µL (blue), 80 ng/µL (purple). (F) Percent of MEG-3 in condensates (Log (I) ≤ 4.5 from histograms in (E) plotted vs. RNA concentration in the presence of increasing salt concentrations assembled as in (D). Each data set represents 8 images collected from 2 experimental replicates. Circles indicate the mean and represent reactions assembled in salt concentrations of 37.5 mM (red), 50 mM (orange), 100 mM (yellow), 150 mM (green), 250 mM (blue), and 500 mM (purple). Bars represent the SD. (G) Percent of MEG-3 in aggregates (Log (I) > 4.5 in histograms in (E) plotted vs. RNA concentration in the presence of increasing salt concentration assembled as in (D). Each data set represents 8 images from 2 experimental replicates. Circles indicate the mean and represent reactions assembled in salt concentrations of 37.5 mM (red), 50 mM (orange), 100 mM (yellow), 150 mM (green), 250 mM (blue), and 500 mM (purple). Bars represent the SD.

**Figure S8.**
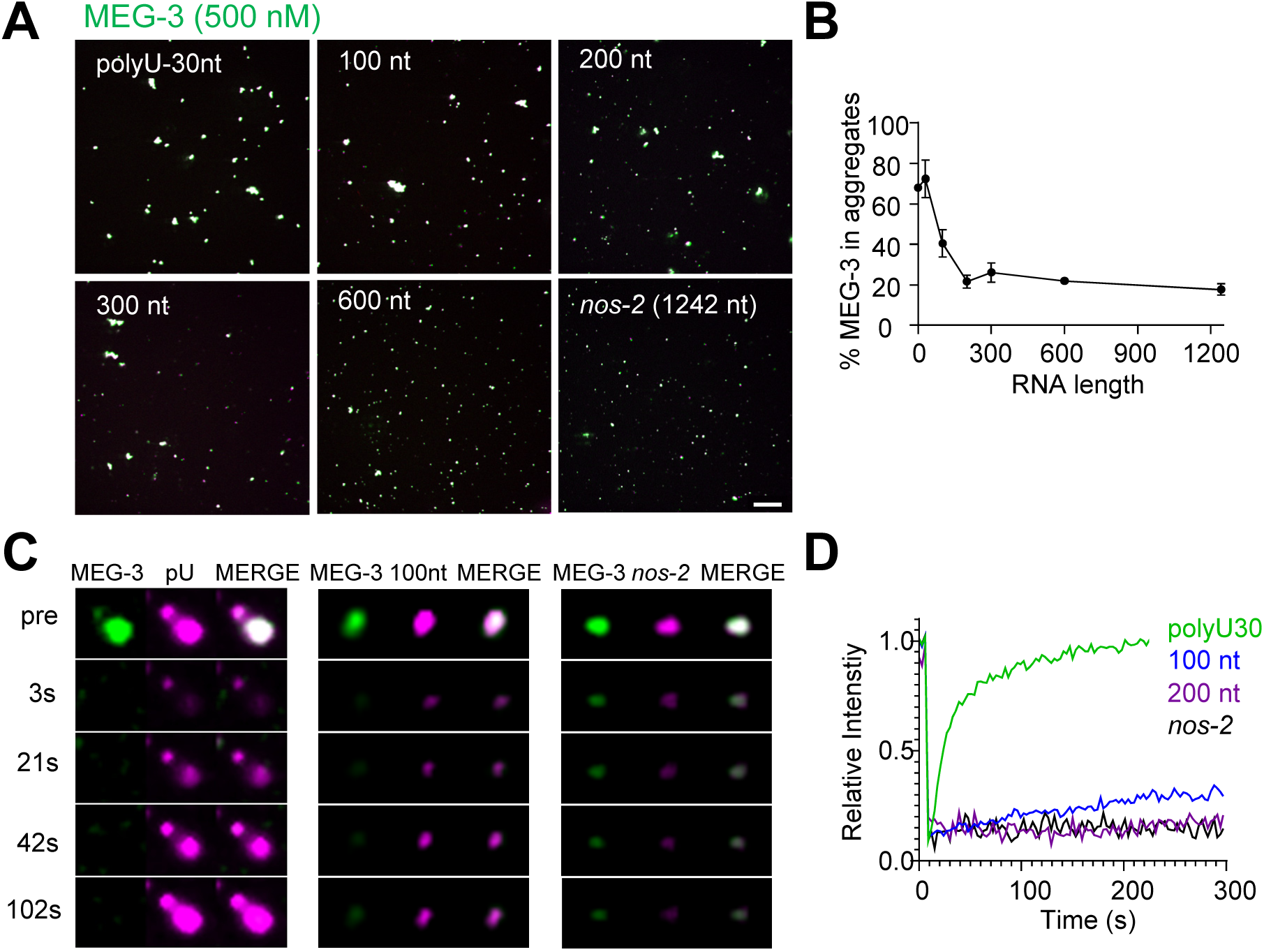
RNA modulates MEG condensation dependent on RNA length. (A) Representative photomicrographs of MEG-3 condensation reactions with indicated RNAs. 100-600 nt RNAs are fragments of *nos-2* (Methods). Reactions contained 500 nM MEG-3 and 20 ng/µL RNA, and salt. MEG-3 (green) was trace labeled with Alexa647 and *nos-2* RNAs (magenta) were trace labeled with Alexa488 or Alexa546, polyU-30nt was trace labeled with fluorescein (Methods). Scale bar is 20 µm. (B) Percent of MEG-3 in aggregates plotted vs. RNA length assembled as in (A). Aggregates were defined as objects with a MEG-3 intensity of Log (I) > 4.6 from histograms in Fig.4E. Each data set includes condensates from 12 images collected from 3 experimental replicates. Circles indicate the mean and bars represent the SD. (Methods). (C) Representative images showing fluorescence recovery after partial photobleaching (FRAP) of condensates assembled as in (A) and incubated for 30 min in condensate buffer. (D) Graph showing fluorescence recovery after photobleaching (FRAP) as described in (C). The RNA intensity of condensates (left panels, n=6) was measured every 3s for 300s before and after bleaching. Values were normalized to initial fluorescence intensity, corrected for photobleaching and plotted as an average. Refer to Fig. 4F recovery rates.

**Figure S9.**
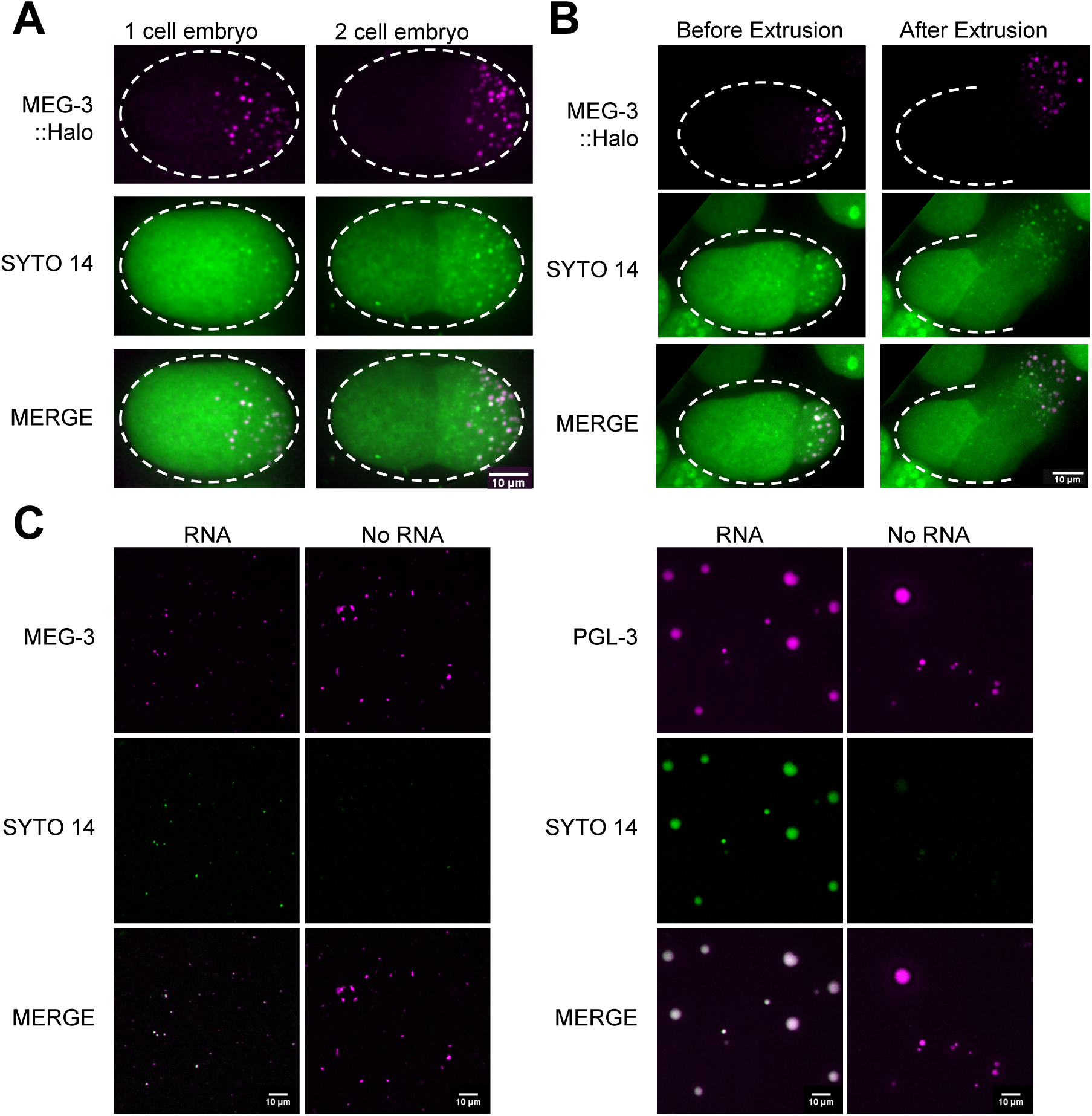
RNA is stably associated with the MEG gel phase *ex vivo*. (A) Photomicrographs of one and two-cell embryos expressing MEG-3::Halo and stained with SYTO 14. Scale bar is 10 μm. (B) Time-lapse photomicrographs of a four-cell embryo expressing MEG-3::Halo and stained with SYTO 14 before and 30s after laser puncture of the eggshell. MEG-3::Halo and SYTO 14 persist in the granule phase. Scale bar is 10 μm. (C) Representative photomicrographs of condensates of MEG-3 or PGL-3 assembled with and without RNA in the presence of SYTO 14 for 30 min. Reactions contained 500 nM MEG-3 or 5 µM PGL-3 and 20 ng/µL *nos-2* (Methods). MEG-3 (magenta) was trace labeled with Alexa647, PGL-3 (magenta) was trace labeled with Alexa647 (Methods). Scale bars are 10 µm.

**Figure S10.**
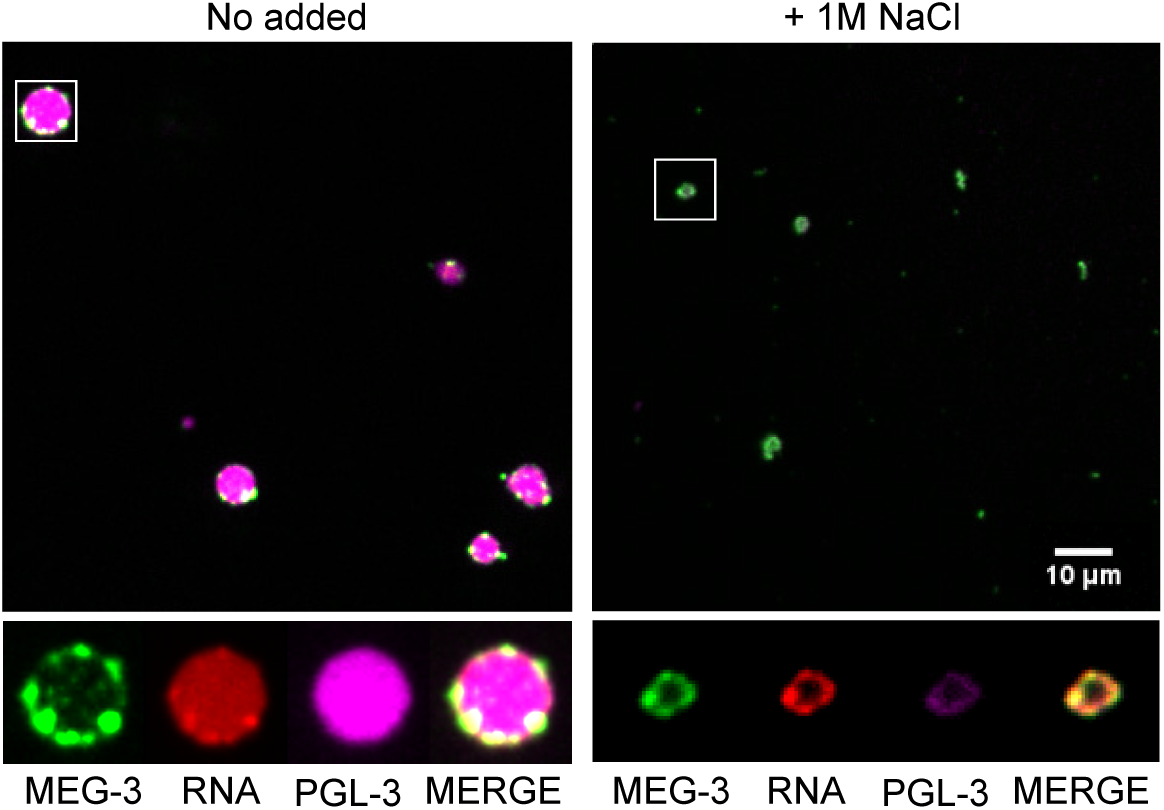
RNA dependent MEG super-assemblies. Additional representative images of condensates assembled as in Fig. 5A.

## Methods

### Worm handling, RNAi, sterility counts

*C. elegans* were cultured according to standard methods(Brenner, 1974).RNAi knockdown experiments were performed by feeding on HT115 bacteria(Timmons and Fire, 1998). Feeding constructs were obtained from Ahringer or OpenBiosystem libraries. The empty pL4440 vector was used as negative control. Bacteria were grown at 37°C in LB + ampicillin (100 µg/mL) media for 5-6 hr, induced with 5 mM IPTG for 30 min, plated on NNGM (nematode nutritional growth media) + ampicillin (100 µg/mL) + IPTG (1 mM) plates, and grown overnight at room temperature. Embryos isolated by bleaching gravid hermaphrodites were put onto RNAi plates directly. To culture larger number of worms for iCLIP and ribosome profiling experiments, worm cultures were started from synchronized L1 (hatched from embryos incubated in M9 overnight) onto NA22 or RNAi bacteria containing plates and growing them to gravid adults. Early embryos were harvested from gravid adults.

To verify the efficiency of RNAi treatments for knocking down *meg* genes, we scored animals exposed to the same RNAi feeding conditions for maternal-effect sterility. For *meg-1(vr10)* strain on *meg-2* RNAi, sterility was 95%±0.6% at 20°C; *meg-1 meg-3 meg-2(RNAi) meg-4(RNAi)* maternal effect sterility was 100% ± 0%. To verify the RNAi efficiency of targeting *mex-3* and *pos-1*, embryonic lethality was assayed. Cohorts of 10-20 mothers were allowed to lay eggs for periods ranging from 2-4 hours. Embryos were then counted, and adults were scored four days later. The embryonic lethality for both *mex-3* and *pos-1* were 100%±0% and 98%±0.2% respectively.

### Strain construction by CRISPR-mediated genome editing

CRISPR generated lines were created as in Paix et al., 2017 as indicated in the Strain Table S3. Guides and repair temples used for CRISPR are listed in Table S4.

### Single molecule fluorescence in situ hybridization (smFISH)

smFISH probes were designed using Biosearch Technologies’s Stellaris Probe Designer. The fluorophores used in this study were Quasar570 and Quasar670. For sample preparation, embryos were extruded from adults on poly-lysine slides (0.01%) and subjected to freeze-crack followed by cold methanol fixation at −20° C. Samples were washed five times in PBS+0.1%Tween20 and fixed in 4% PFA (Electron Microscopy Science, No.15714) in PBS for one hour at room temperature. Samples were again washed four times in PBS+0.1%Tween20, twice in 2x SCC, and once in wash buffer (10% formamide, 2x SCC) before blocking in hybridization buffer (10% formamide, 2x SCC, 200 ug/mL BSA, 2mM Ribonucleoside Vanadyl Complex, 0.2 mg/mL yeast total RNA, 10% dextran sulfate) for 30 minutes at 37° C. Hybridization was then conducted by incubating samples with 50-100 nM probe solutions diluted in hybridization buffer overnight at 37° C. Following hybridization, samples were washed twice in wash buffer at 37° C, twice in 2x SCC, once in PBS-Tween20 (0.1%), and twice in PBS. Lastly, samples were mounted using VECTASHIELD Antifade Mounting Media with DAPI or Prolong Diamond Antifade Mountant.

### Immunostaining

As in sample preparation for smFISH experiments, embryos were extruded from adults and subjected to freeze-crack on poly-lysine slides followed by cold methanol fixation for 15 minutes and then cold acetone for 10 minutes. Slides were blocked twice in PBS-Tween20 (0.1%)-BSA (0.1%) for 30 minutes at room temperature, and incubated with 90 μl primary antibody overnight at 4°C in a humidity chamber. Antibody dilutions (in PBST/BSA): mouse K76 1:10 (DSHB), Rat α-OLLAS 1:80 (Gift from Dr. Jeremy Nathans), mouse α-FLAG M2 1:500 (Sigma F3165). Secondary antibodies (Molecular Probes/Thermo Fisher Sci.) were applied for 1∼2 hours at room temperature.

### Confocal microscopy

Fluorescence confocal microscopy was performed using a Zeiss Axio Imager with a Yokogawa spinning-disc confocal scanner. Embryo images were taken using Slidebook v6.0 software (Intelligent Imaging Innovations) using a 63x objective. Embryos were staged by DAPI-stained nuclei in optical Z-sections and multiple Z-sections were taken to include germ cells. For *In vitro* condensation reactions, images are single planes taken using a 40x objective unless otherwise indicated. For fluorescence super-resolution microscopy, images were acquired using ZEISS LSM 880-AiryScan (Carl Zeiss) equipped with a 63X objective. Images were processed using ZEN imaging software (Carl Zeiss). Equally normalized images were exported via either Slidebook v6.0 or ZEN, and contrasts of images were equally adjusted between control and experimental sets. For *in vitro* fluorescence recovery after photobleaching experiments, images were acquired using Zeiss LSM 800 GaAsp. Images are single confocal planes imaged using a 63x objective every 3s during a recovery phase of 300s. All image analyses were conducted using the Fiji image-processing package (http://fiji.sc/Fiji).

### Quantification of RNA granule size

For measurements reported in Fig. 1D, 9 Z-planes (0.8 μm step size) from the center of an embryo were extracted for image analysis. To identify RNA granules, minimum and maximum values for thresholding were set as follows: minimum= the mean intensity of the background signal plus one standard deviation of the background intensity; maximum= the mean intensity of the background signal plus 6 standard deviations of the background intensity. After thresholding, the nucleus counter cookbook plugin in FIJI was used to identify RNA granules in germ cells. Objects of less than 2 pixels were filtered out to minimize noise, a watershed filter was applied to improve separation of granule signals close in proximity, and the image was converted to a binary image by the “Current” method. Measurements for granule area were extracted from the ROI manger. For Fig. 1C, mean and standard deviation of granule size from at least 4 embryos were plotted against average read counts from two MEG-3::GFP iCLIP experiments.

### Temperature shifts

Temperature shift experiments were performed by transferring gravid worms grown at 20°C to pre-warmed 30°C plates for 15 minutes. After heat shock, worms were immediately dissected for smFISH experiments or live imaging. To quantify the ratio of smFISH signal in MEG-3::GFP granules, 8 single Z planes were extracted and used for image analysis as shown in Figure 2E.

MEG-3::GFP granules were identified using the nucleus counter plugin as described above. The minimum threshold was set to the 2 times the mean intensity of the background signal of the image; and the maximum threshold was calculated by adding 6 standard deviations of the background intensity. A mask generated from objects identified by the nucleus counter plugin was applied to the raw image to extract RNA or GFP intensity. To remove background signal, the mean intensity of an object across the nucleus was measured, and subtracted from calculated RNA or GFP intensities. To calculate the ratio of RNA signal/ MEG-3::GFP granule signal in each selected Z plane, the sum of intensities of RNA in identified objects (*I*_smFISHg_) was divided by the total intensity of the MEG-3::GFP in the same objects (*I_meg3g_*). The intensity ratio of RNA in MEG-3 granules (*I*_smFISHg_/ *I_meg3g_*) before and after heat shock was compared. Each data point represents data from one Z plane acquired from two embryos (8 planes per embryo).

#### Translation inhibitor treatments

For drug treatment, *ptr-2* RNAi was used to permeabilize the egg shell. A 12.5 mg/mL (20X) Puromycin stock solution (sigma, P8833) was made with osmolarity calibrated Egg buffer [118 mM NaCl, 48 mM KCl, 2 mM CaCl_2_, 2 mM MgCl_2_ 25 mM HEPES pH 7.3, 340±5 mOsm]. A 50 mg/mL (200X) Cycloheximide stock solution was made in ethanol. RNAi treated gravid worms were dissected and permeabilized embryos were released into drug containing egg buffer for 1 hour in a humidity chamber to maintain vapor pressure. After drug treatment, excess buffer was removed and embryos were subjected to image acquisition as described above using nucleus counter cookbook plugin. Both puromycin and cycloheximide induced cell cycle arrest.

#### RNA enrichment in germ granule vs cytosol

smFISH quantification was conducted using Imaris Image Analysis Software visualization in 3D space. The boundary/volume for the germ cell cytosol and germ granules was created by Surface function using MEG-3::GFP signal in Imaris. The sum of intensity of the germ cell cytosol and granules were extracted and the percentage of RNA enrichment in germ granules was calculated.

### Protease inhibitor mix for Immunoprecipitation

We prepared freshly made 100x protease inhibitor mixes for the immunoprecipitation step in the iCLIP protocol. Protease inhibitor mix1(100x): Antipain (3 mg), Leupeptin(5 mg), Benzamidine(10 mg), AEBSF(25 mg) and phosphoramidon (1 mg) were resuspended in 1 mL Phosphate-buffered saline (PBS). Protease inhibitor mix2 (100x): 500 µL of 10 mg/mL Aprotinin, 400 µL of 10 mM Besttatin, 100 µL of 10 mM E64, and 100 µL of 10 mg/mL Trypsin inhibitor were mixed together in H2O.

### iCLIP and library preparation

#### Crosslinking, immunoprecipitation and nuclease treatments

The iCLIP protocol was adapted from Huppertz, *et al*. (Huppertz et al., 2014) with some modifications as detailed below. *C. elegans* embryos collected from ∼6.7×10^7^ gravid adults were seeded on 10 cm petri dishes and irradiated 3 times with 400 mJ/cm^2^ at UV 254 nm by Stratalinker. Crosslinked embryos were collected and resuspended in Immunoprecipitation buffer (IP buffer) with freshly made protease inhibitors as described in this section [300 mM KCl, 50 mM HEPES pH7.4, 1 mM EGTA, 1 mM MgCl_2_, 10% glycerol, 0.1% NP-40]. Samples were lysed in a Spex 6870 freezer mill followed by centrifugation at 4°C 21,000xg for 30 minutes to remove embryo debris. Cleared lysates were subjected to immunoprecipitation using anti-GFP antibody (Roche 11814460001) conjugated to protein G magnetic beads (Thermofisher) in the presence of 200unit of RNaseOut per milliliter of lysates. Immunoprecipitated fractions were then washed extensively with IP buffer, low salt buffer [150 mM NaCl, 50 mM HEPES pH7.4, 0.1% SDS, 0.5% NP-40, 1 mM MgCl_2_, 1 mM EGTA] and high salt buffer [500 mM NaCl, 50 mM HEPES pH7.4, 0.1% SDS, 0.5% NP-40, 1 mM MgCl_2_, 1 mM EGTA]. Bound fractions were then washed with 1xPBS to remove excess salt followed by RQ1 DNase treatment at 37° C for 10 minutes [4 units of RQ1 (Promega M6101) and 60 units of SUPERaseIN (ThermoFisher AM2694)]. To perform partial RNA digestion, RQ1 DNase treated bound fractions were washed with 1 mL low salt buffer, 1 mL high salt buffer, 1 mL 1x PNK buffer [50 mM Tris-HCl pH7.4, 10 mM MgCl_2_, 0.5% NP-40] and finally resuspend in 500 µL MNase reaction buffer [NEB, M0247, 1 unit/mL in 1x reaction solution]. The MNase reaction was immediately transferred to a thermomixer for 2 minutes at 37°C. The MNase reaction was stopped with ice-cold 1x PNK buffer with 5 mM EGTA followed by 2x 1 mL high salt buffer, 2x 1 mL low salt buffer and 1x 1 mL PNK buffer.

#### L3 Adapter ligation

A dephosphorylation step is necessary to remove 3’ end phosphates that prevent adapter ligation. Beads were resuspended in 20 µL of dephosphorylation reaction [4 µL of 5x PNK reaction buffer, 0.5 µL T4 PNK (NEB M0201), 0.5 µL RNasin (Promega N2111), 15 µL H_2_O] at 37° C for 20 minutes followed by washes with PNK buffer. Beads were resuspended in in 20 µL ligation mixture [2 µL of 10x RNA ligation buffer, 2 µL of T4 RNA ligase II truncated KQ (NEB M0373), 0.5 µL of SUPERaseIN, 1.5 µL of 20 µM pre-adenylated L3-App adapter, 4 µL of PEG8000, 10 µL H_2_O] at 16° C for overnight.

#### 5’ end labeling, SDS-PAGE and nitrocellulose transfer

Ligated RNA samples were washed twice with high salt buffer and twice with PNK buffer. Supernatants were removed and samples were resuspended in 80 µL of hot PNK mix for 5’ end labeling [8 µL of 10x PNK buffer, 1 µL of P^32^ ATP, 4 µL of T4 PNK, 67 µL H_2_O] at 37° C for 10 minutes. Remove unlabeled hot ATP by wash beads with wash buffer [20 mM Tris-HCl pH7.4, 10 mM MgCl_2_, 0.2% Tween-20]. Samples were loaded on a 4-20% TGX protein gel (Bio-Rad 4561093) and transferred to a nitrocellulose membrane.

#### RNA isolation, Reverse transcription, cDNA circularization and PCR amplification

These procedures were performed as described in Huppertz, *et al*. (Huppertz et al., 2014).

### RNA extraction and preparation of mRNA-seq library preparation

RNA was extracted from embryos or cleared embryo lysates using TRIZOL. The aqueous phase was transferred to Zymo-SpinTM IC Column (Zymo research R1013) for concentration and DNase I treatment as described in manual. RNA quality was assayed by Agilent Bioanalyzer using Agilent RNA 6000 Pico Chip. All RNAs used for library preparation had RIN (RNA integrity number) >9. For mRNA-seq library construction, 0.5 µg of total RNA was treated with Ribo-Zero Gold Epidemiology rRNA Removal Kit. Libraries were then prepared following the TruSeq RNA Library Prep Kit v2 instruction. All sequencing was performed using the Illumina HiSeq2500 at the Johns Hopkins University School of Medicine Genetic Resources Core Facility.

### Preparation of libraries for ribosome profiling

Synchronized L1 worms were seeded on plates containing HT115 bacteria transformed with pL4440 vector and cultured at 25°C for ∼48 hours. Additional bacteria were added to ensure enough food to support development. Early embryos were collected by bleaching gravid hermaphrodites. Small aliquots of embryos were collected from each experiment and staged by DAPI-staining. 70%±7% of embryos were before or at ∼100 cell stage. ∼400 μL packed embryos were then resuspended in 2 mL footprint lysis buffer [20 mM Tris-Cl (pH8.0), 140 mM KCl, 1.5 mM MgCl2, 1% Triton X-100, 0.1 mg/mL CHX] and lysed in a Spex 6870 freezer mill. After clarification lysate by sequential centrifugation at 3000rpm followed by 17,000xg, 100 µL (5% of lysate) of lysates were saved for mRNAseq. For ribosome profiling, lysates containing 300 μg of total RNA were treated with 100 units of RNaseI (Ambion) for 30 min at 25°C. 40 units of SUPERaseIN (ThermoFisher) were added to prevent further digestion from RNaseI. Monosomes were isolated by sucrose gradients (10-50%) and centrifuged at 40,000 rpm for 3 hours in a SW41 rotor (Beckman Coulter). The extracted RNA was size-selected from 15% denaturing PAGE gels, cutting between 15-34 nt. An oligonucleotide adapter was ligated to the 3’ end of isolated fragments. After ribosomal RNA depletion using RiboZero (Illumina), reverse transcription using SuperScript III reverse transcriptase (Thermo Fisher Scientific), circularization using CirLigase I (Lugicen) and PCR amplification(Schuller et al., 2017). Libraries were sequenced on a HiSeq2500 machine at facilities at the Johns Hopkins Institute of Genetic Medicine.

### High-throughput sequencing analysis

#### mRNA sequencing

sequencing reads were aligned to the UCSC ce10 *C. elegans* reference genome using HISAT2(Kim et al., 2015). Reads aligning to genetic features were then counted using HTSeq-count(Anders et al., 2015) and analyzed for differential expression analysis using DESeq2(Love et al., 2014). For analysis shown in Fig. 3C, differential expression analysis was done using Tophat (V.2.0.8) and Cufflink (V.2.0.2). The command lines for Tuxedo suit are listed as below:

For each biological sample, sequencing reads were first mapped to ce10 reference genome using tophat2:

$ tophat2 -p 12 -g 1 --output-dir <output> --segment-length 20 --min-intron-length 10
--max-intron-length 25000 -G <gene.gtf> --transcriptome-index <Name.fastq>

For differential gene expression analysis, sets of independent mutant and control mapped reads (e.g biological replicates) were used in cuffdiff analysis:

$ cuffdiff -p 12 -o <output> --compatible-hits-norm --upper-quartile-norm -b
<genome.fa> <genes.gtf> <tophat output_sample 1, tophat output_sample 2, tophat
output_sample 3,..> <tophat output_control1, tophat output_control2, tophat
output_control3,.. >

#### Ribosome profiling

Libraries for WT and *meg-3/4* embryos were trimmed to remove the ligated 3’ linker (CTGTAGGCACCATCAAT) with skewer(Jiang et al., 2014). For the rest of our libraries, the 3’ adapter (NNNNNNCACTCGGGCACCAAGGA) was trimmed, and 4 random nucleotides included in the RT primer (RNNNAGATCGGAAGAGCGTCGTGTAGGGAAAGAGTGTAGATCTCGGTGGTCGC/iSP18/TTCAGACGTGTGCTCTTCCGATCTGTCCTTGGTGCCCGAGTG) were removed from the 5’ end of reads. Trimmed reads longer than 15 nt were aligned to worm ribosomal and non-coding RNA sequences using STAR(Dobin et al., 2013) with ‘--outFilterMismatchNoverLmax 0.3’. Unmapped reads were then mapped to genome using the following options ‘—outFilterIntronMotifs RemoveNoncanonicalUnannotated --outFilterMultimapNmax 1 --outFilterMismatchNoverLmax 0.1’. Aligned reads were than counted and analyzed using HTseq-count(Anders et al., 2015), DEseq2(Love et al., 2014) and custom R code (RStudio 1.2)

#### iCLIP data analysis

The 5’ barcodes (NNN-4 nt indexes – NN) and 3’ adaptor (AGATCGGAAG) for iCLIP library construction were listed in Huppertz, *et al*. (Huppertz et al., 2014). iCLIP sequencing reads were trimmed to remove 3’ adaptor and 5’ randomized barcodes using - fastx_clipper and custom python codes base on Ule lab GitHub depository (https://github.com/jernejule/non-coinciding_cDNA_starts). Trimmed reads were aligned to *C. elegans* ws235 reference genome using HISAT2 (Kim et al.,2015), and PCR duplicated reads were removed. To determine the distribution of mapped read across genome, an R package RNA centric annotation system (RCAS) was used to generate the plot shown in Fig. S4A. Reads aligning to genetic features were then counted using HTseq-count. This information was used to plot Fig. 1A, 1B, 1D, and Fig. 2A. To generate the list of “MEG-3 bound transcripts”, genes identified in both MEG-3::GFP iCLIP experiments were kept and filtered out genes with read counts fewer than 60 (background level base on GFP iCLIP shown in Fig.1A), which yielded a list of 657 “MEG-3 bound transcripts”.

To identify potential MEG-3 binding motifs, MEG-3::GFP and control GFP iCLIP mapped reads were used in peak caller PEAKachu in galaxy server (https://usegalaxy.eu/) with options – Menimum cluster Expression Fraction 0.01 --Minimum Block Overlap 0.5 –Minumum Block Expression 0.1 –Mad Multiplier 2.0 –Fold Change Threshold 1.5 – Adjusted p-value Threshold 0.1. Identified peaks with additional 15 nt extensions from both ends were used by MEME suite to search for sequence motif, and no motif reached the E-value cut off (E-value)<1x 10-5. An additional iCount analysis package (Curk et al. (2019), https://icount.readthedocs.io/en/latest/index.htmL) with the same options described in the tutorial document (https://icount.readthedocs.io/en/latest/ref_CLI.html) was used to identify significant peaks. 15 nt extensions were added to both sides of peaks and followed by MEME suite motif analysis. No motif with an E-value < 1×10-5 was found. Therefore, we conclude that MEG-3 binds RNAs without any sequence bias.

To plot the MEG-3 binding profile, bamCompare and computeMatrix in the deepTools package (http://deeptools.readthedocs.io/en/latest/) were used to compute mapped read coverage. Command lines were listed as below:

$ bamCoverage -b <inpit> -o <output.bw> -bs 1 -p 8 -ignore chrM --exactScaling -- smoothLength 3
$ computeMatrix scale-regions -S <input.bw> -R <MEG-3-bound transcripts.bed> -b 800-a 1500 -m 2000 -bs 1 --skipZeros --sortUsing max -p 8 -o <output.gz>
$ plotProfile -m <input.gz> --out <output.pdf> --colors blue green --perGroup

### Gene list: MEG-3 bound and P granule transcripts

Read counts obtained in the control GFP iCLIP were used to define background threshold (read count = 60.) We defined MEG-3 bound transcripts by excluding transcripts with MEG-3::GFP iCLIP read counts < 60 as shown in Fig. 1A (stippled horizontal line at log_e_60 = 4.1). To define P granule transcripts, we used the rank order of *F35G2.1* (Rank 388), the left most gene in the *nos-2* cluster as shown in Fig. 1C, as the cut off. Genes with rank order better than 388 in either one of MEG-3::GFP iCLIP were defined as P granule transcripts (Table S1).

### Protein purification and labeling

#### Purification of MEG-3 His-tagged fusion

MEG-3 full-length (aa1-862) fused to an N-terminal 6XHis tag in pET28a was expressed and purified from inclusion bodies using a denaturing protocol(Smith et al., 2016). MEG-3 was grown in Rosetta (DE3) cells at 37°C in terrific broth + ampicillin (100 µg/mL) to an OD600 of ∼1.0 and induced with 1 mM IPTG at 16° C for 16 hr. Cells were resuspended in Buffer A (20 mM HEPES pH 8.0, 1000 mM KCl, 10% (vol/vol) glycerol, 0.5% Triton-X100, 2 mM DTT, 0.4 mM PMSF, and Roche proteinase inhibitors), lysed by sonication, and spun at 13,000 rpm for 30 min. The supernatant was discarded and the pellet containing MEG-3 inclusion bodies was resuspended in Buffer A, briefly sonificated, and spun at 13,000 rpm. The pellet was solubilized overnight at 4°C in Buffer A with 6 M Urea. The solubilized protein was filtered (0.45 µm), and passed over a HisTRAP 5 mL column (GE Healthcare). Bound protein was washed with Buffer B (20 mM HEPES pH 8, 1 M KCl, 25 mM Imidazole pH 6, 10% (vol/vol) glycerol, 6 M urea, 2 mM DTT) and eluted in Buffer C (20 mM HEPES pH 7.4, 1 M KCl, 250 mM Imidazole pH 6, 10% (vol/vol) glycerol, 6 M urea, 2 mM DTT). Protein containing fractions were concentrated to 3 mL and further purified by size exclusion using a HiPrep 16/60 Sephacryl S-500 HR (GE Healthcare) in Buffer D (20 mM HEPES pH 7.4, 1 M KCl, 10% (vol/vol) glycerol, 6 M urea, 2 mM DTT). Aliquots of peak elution fractions were run on 4–12% Bis Tris gels, and stained with Simply Blue Safe Stain (ThermoFisher Waltham, MA). Protein was concentrated to a final concentration of 2-5 mg/mL, aliquoted, snap frozen in liquid nitrogen, and stored at −80°C.

Purification of MBP-TEV-PGL-3 was expressed and purified as described(Putnam et al., 2019) with the following modifications: MBP was cleaved using homemade TEV protease instead of commercial. A plasmid expressing 8X-His-TEV-8X-Arg tag protease was obtained from Addgene and purified according to the published protocol(Tropea et al., 2009). Before loading cleaved PGL-3 protein on to a heparin affinity matrix, cleaved MBP-6X-His and 8X-His-TEV protease were removed using a HisTRAP column (GE Healthcare).

#### Protein labeling

Proteins were labeled with succinimidyl ester reactive fluorophores from Molecular Probes (Alexa Fluor™ 647, Alexa Fluor^TM^ 555, or DyLight™ 488 NHS Ester) following manufacturer instructions. Free fluorophore was eliminated by passage through three Zeba™ Spin Desalting Columns (7K MWCO, 0.5 mL) into protein storage buffer. The concentration of fluorophore-labeled protein was determined using fluorophore extinction coefficients measured on a Nanodrop ND-1000 spectrophotometer. Labeling reactions resulted in ∼ 0.25-1 label per protein. Aliquots were snap frozen and stored. In phase separation experiments, fluorophore-labeled protein was mixed with unlabeled protein for final reaction concentrations of 25-100 nM of fluorophore labeled protein.

### *In vitro* RNA preparation

mRNAs were transcribed using T7 or SP6 mMessageMachine (Thermofisher) using manufacturer’s recommendation. 1 µL of chromatide ChromaTide™ Alexa Fluor™ 488-5-UTP or 546-14-UTP (Thermofisher) were added to transcription reactions to fluorescently trace label mRNAs. Template DNA for transcription reactions was obtained by PCR amplification from plasmids. *nos-2* fragments were generated by PCR amplification from the 5’ end of the full length *nos-2* template DNA. Free NTPs and protein were removed by lithium chloride precipitation. RNAs were resuspended in water and stored at −20°C. The integrity of RNA products was verified by agarose gel electrophoresis.

30 nt oligo RNAs were ordered from IDT either unlabeled or with a 3’ FAM modification. Oligos were resuspended in water aliquoted and stored at −80°C. Labeled and unlabeled oligo RNAs were mixed together and used at final concentrations of 20 ng/uL including 25 nM fluorescently labeled oligo. Oligo sequences: polyU-30 nt,

Oligo1:UCUUUUUCCUCUACACAACUUUUUUUUACA,

Oligo2:UCUAGCAUGUCGGCCCUUGAUCGGCUGACA,

Oligo3:CAUCCUCAAAUCUACGAUCAACAGAUGCAG,

Oligo4:AUGGGUGUGAUGCCACAUACGCAGCCGGCA

### *In vitro* condensation experiments and analysis

Protein condensation was induced by diluting proteins out of storage buffer into condensation buffer containing 25 mM HEPES (pH 7.4), salt adjusted to a final concentration of 150 mM (37.5 mM KCl, 112.5 mM NaCl), and RNA. Unless otherwise indicated, for all co-assembly experiments, we used 500 nM MEG-3, 5 μM PGL-3/PGL-3 and 20 ng/µL RNA. MEG-3 and PGL-3 solutions contained 25 nM fluorescent trace labels with either 488, 555, or 647 (indicated in figure legends). Condensate reactions with the RNA dye contained a final concentration of 100 nM SYTO 14. Condensation reactions were incubated at room temperature for 30 min or as indicated, before spotting onto thin chambered glass slides (ERIE SCIENTIFIC COMPANY 30-2066A) with a coverslip. Images used for quantification are single planes acquired using a 40x oil objective over an area spanning 171 x 171 μm.

To quantify the ratio of protein or RNA in condensates, a mask was created by thresholding images, filtering out objects of less than 4 pixels to minimize noise, applying a watershed filter to improve separation of objects close in proximity, and converting to a binary image by the Otsu method using the nucleus counter cookbook plugin. Minimum thresholds were set to the mean intensity of the background signal of the image plus 1-2 standard deviations. The maximum threshold was calculated by adding 3-4 times the standard deviation of the background. Using generated masks, the integrated intensity within each object was calculated. To remove non-specific background signal the mean intensity of an image field in the absence of the labeled component was subtracted from each pixel yielding the total intensity of each object.

Histograms of MEG-3 intensity were generated by taking the log(10) of total intensity for each MEG-3 object, Log(I). Objects in 2-3 experimental replicates of 4 images were identified and quantified as described above The number of objects for each Log(I) value was binned (bin size = 0.2 Log(I) units), and normalized to the total number of objects. The percent of objects in each bin was averaged for each experimental replicate.

The percent of MEG-3 in aggregates or condensates was determined by comparing histograms of reactions in which all objects are aggregates (500 mM NaCl or no RNA) or condensates (150 mM NaCl and 20-80 ng/µL *nos-2* RNA) as illustrated in Fig. 4B,E and Fig. S7E. The minimum at the intersection of the two conditions was calculated. The percentage of MEG-3 objects with an intensity above or equal to the intersection were classified as aggregates and the fraction of objects with an intensity below the intersection were classified as condensates. To calculate the fraction of RNA in MEG-3 condensates/aggregates, the background corrected sum of RNA fluorescent intensity in all MEG-3 objects was divided by the total intensity of RNA fluorescence in the imaged area.

Radii of MEG-3 condensates were estimated by imaging condensation reactions of 500 nM MEG-3 and 40 ng/µL RNA. Images used for quantification were single planes acquired using a 100x oil objective over an area spanning 68 x 68 μm. 4 experimental replicates of 16 images were identified and quantified (>1500 objects/replicate) as described above, and radii were calculated from the area of each object. Calculated radii are an overestimate and represent upper limits for actual condensate size. The number of objects for each radii was binned (bin size = 0.06 µm), and normalized to the total number of objects. The percent of objects in each bin was averaged for each experimental replicate.

### Fluorescence Recovery after Photobleaching (FRAP)

20 µL condensation reactions (prepared as described above) were added to a chambered coverglass (Grace BioLabs) and imaged using a Zeiss LSM 800 GaAsp. Bleaching was performed using 100% laser power in the 488, 546, or 647 channels. Regions slightly larger than the condensates (radius ≈ 3 µM) were photobleached. A single confocal plane was imaged using a 63x objective every 3s during a recovery phase of 300s.

FRAP analysis was performed as described in Putnam *et al.*, 2019(Putnam et al., 2019). Briefly, fluorescence recovery was corrected for background and normalized to the initial granule intensity using the equation: n*I* = (*I-I^bkg^*)/(*I^i^*-*I^bkgi^*), where n*I* is the background corrected and normalized fluorescence intensity, *I* is the intensity of the FRAPed granule, *I^bkg^* is the fluorescence intensity outside of the condensate, *I^i^* is the initial intensity before bleaching, and *I^bkgi^* is the initial background intensity. Recovery rates were determined by fitting individual traces to a first order equation n*I* = (A^rec^·(1-e*^-**k**^***^t^**), where A^rec^ is the fluorescence recovery amplitude and ***k*** is the rate of fluorescence recovery.

### *Ex vivo* Extrusion Experiments

1 mM SYTO 14 RNA Dye (ThermoFisher) and 1 mM JF_646_ (JF_646_, Janelia(Grimm et al., 2015)) dissolved in DMSO were diluted 500 fold into osmolarity calibrated Egg buffer [118 mM NaCl, 48 mM KCl, 2 mM CaCl_2_, 2 mM MgCl_2_ 25 mM HEPES pH 7.3, 340±5 mOsm] to reach 2 µM stock solutions. *perm-1(*RNAi) adult gravid hermaphrodites expressing MEG-3::HALO(Wu et al., 2019) were dissected and egg shell permeabilized embryos were released into 10 µL egg buffer containing 1 µM SYTO 14 and JF_646_ for 10 min in a humid chamber to prevent evaporation. After drug treatment, embryos were washed 3X with egg buffer without drug. Approximately 200-20 µm polysterene beads (Bangs Laboratories) suspended in egg buffer were added to prevent embryo compression, and placed on slides for imaging. Embryo contents were extruded by puncturing the eggshell near the anterior region of the germline blastomere using a 3i Ablate!^TM^ laser system at 532 nm pulse setting with a power level of 155(Putnam et al., 2019). All embryo images are Z stack maximum projections using a Z step size of 1 μm, spanning the depth of the embryo. Images were acquired in the 488 and 647 channel every 10s using a 63x objective.

To quantify SYTO 14 and MEG-3::Halo persistence in granules, photomicrographs acquired as described above were analyzed using FIJI. Not all SYTO 14 granules were MEG-3::Halo postive, and are potentially P bodies (gallo). Only SYTO 14 granules also positive for MEG-3::Halo were quantified. MEG-3::Halo granules were identified using the nucleus counter plugin as described above. The total intensity of objects was quantified for both 488 (SYTO 14) and 647 (Halo) channels. Total fluorescence intensity was calculated before (I_B_) and after (I_A_) extrusion and used to calculate a fluorescence ratio (I_A_/I_B_). Photobleaching was minimal for MEG-3::Halo; however it was significant for SYTO 14. To correct for photobleaching, total fluorescence intensity was corrected for photobleaching using the average photobleaching rate calculated from the cytoplasm of intact embryos in the imaging area. For some embryos, granules left the field of view and could not be counted. The I_A_ /I_B_, therefore, is a minimal estimate of the fraction MEG-3 and SYTO 14 that remained in the granule phase after extrusion.

### Graphing and data fitting

All data was plotted and statistical analysis was conducted using Graphpad Prism 7 software. Fitting of recovery curves in FRAP experiments was conducted using Kaleidagraph (Synergy) software.

### Data and Code Availability

Sequencing data has been deposited onto the Gene Expression Omnibus (GEO) and can be found using the following accession numbers:

xxxxxxxxxxxxx

xxxxxxxxxxxxx

xxxxxxxxxxxxx

