## Supplemental Table 3 for "Recruitment of mRNAs to P granules by gelation with intrinsically-disordered proteins"

| Supplementary Table 1: List of strains used in this study | | | | | |
|---|---|---|---|---|---|
| Strain | Genotype | Description | Method | Derived | Citation |
| JH3503 | *meg-3(ax3054)* X | MEG-3::meGFP | CRISPR/Cas9 genome editing | N2 | Smith, *et al.* 2016 |
| JH3269 | *pgl-1(ax3122)* IV | PGL-1::GFP | CRISPR/Cas9 genome editing | N2 | Putnam, *et al.* 2019 |
| JH3193 | *nos-2(ax2049[3xFLAG::nos-2]) II* | 3XFLAG ::NOS-2 | CRISPR/Cas9 genome editing | N2 | Paix, *et al* 2014 |
| JH3605 | *Y51F10.2(ax4319[Y51F10.2:: OLLAS]) I* | Y51F10.2::OLLAS | CRISPR/Cas9 genome editing | N2 | This study |
| EGD364 | *meg-3(egx4)* X | MEG-3::Halo | CRISPR/Cas9 genome editing | N2 | Wu, *et al.* 2019 |
| JH3475 | *meg-3(ax3055)* X *meg-4(ax3052)* X | *meg-3*(Δ) *meg-4*(Δ) | CRISPR/Cas9 genome editing | JH3477 | Smith, *et al.* 2016 |
| WM527 | *prg-1(ne4523 [gfp::tev::flag::prg-1]) I* | GFP::TEV::FLAG:: PRG-1 | CRISPR/Cas9 genome editing | N2 | Shen *et al.*, 2018 |
| JH3357 | *nos-2(ax3103)* | *nos-2* mutant | CRISPR/Cas9 genome editing | N2 | Lee, *et al* 2017 |
| SS608 | pgl-3(bn103) | *pgl-3*(Δ) | | | Karashima, *et al.* 2004 |
| SX922 | *prg-1(n4357)I* | *prg-1*(Δ) | | | Caenorhabditis Genetics Center |
| JH3229 | *meg-1(vr10) X meg-3(tm4259)X* | *meg-1 meg-3* mutant | | | Wang *et al.* 2014 |
| JH2878 | *meg-1(vr10)* | *meg-1* mutant | | | Leacock and Reinke. 2007 |
| RB1413 | *Y51F10.2(ok161)* I | *Y51F10.2* mutant | | | Caenorhabditis Genetics Center |

meGFP: monomeric enhanced GFP (A206K)
